# Probing protein-protein and protein-substrate interactions in the dynamic membrane-associated ternary complex of Cytochromes P450, b5, and Reductase

**DOI:** 10.1101/707729

**Authors:** Katherine A. Gentry, G. M. Anantharamaiah, Ayyalusamy Ramamoorthy

## Abstract

Cytochrome P450 (cytP450) interacts with two redox partners, cytP450 reductase and cytochrome-b_5_, to metabolize substrates. Using NMR, we reveal changes in the dynamic interplay when all three proteins are incorporated into lipids nanodiscs in the absence and presence of substrates.

---

Cytochrome P450s (cytP450) are a ubiquitous superfamily of enzymes responsible for the metabolism of a variety of compounds from vitamins to fatty acids to over 70% of the drugs on the pharmaceutical market.^[1–5]^ Per turn of its catalytic cycle which requires two electrons, cytP450 inserts a molecule of activated oxygen into a hydrophobic substrate. Both electrons can be provided by cytochrome P450 reductase (CPR), cytP450’s obligate redox partner, or the second can be donated by cytochrome b_5_ (cytb_5_).^[6]^

CPR is an 80 kDa protein that consists of the FAD/NADPH binding domain, the FMN binding domain (FBD), a linker region connecting the two flavin domains, and an N-terminal transmembrane domain.^[5, 7]^ After being reduced by NADPH, electrons flow from the FAD binding domain to FMN in the FBD which then directly donates the electrons to the heme in the active site of cytP450. In this study, we utilize a truncated version of CPR, the full-length FBD (flFBD), consisting of only the FBD along with the N-terminal transmembrane domain. This flFBD domain has been used previously as the minimal necessary domain of CPR to interact with cytP450 ^[8–10]^.

Cytb_5_, the third protein of this ternary complex, is a ~15.7 kDa protein only capable of donating the second electron to cytP450 because of the disparity in redox potentials between it and ferric cytP450.^[5, 11, 12]^ Cytb_5_ has been shown to increase, decrease, or do nothing to cytP450’s metabolism depending on the isoform of cytP450 and the type of substrate involved.^[2, 13, 14]^ In some cases, such as that of cytP450 17A1, cytb_5_ is known to favor a specific reaction and product formation.^[15, 16, 17]^ Due to the mystery still surrounding cytb_5_’s functional properties, we further investigated cytb_5_‘s role in regulating cytP450 drug metabolism.

All three proteins, cytP450, cytb_5_, and flFBD, contain single transmembrane helices that anchor them to the membrane (Figure 1A). In cytP450’s case, even the globular, ‘soluble’ domain, specifically the F/G-loop, interacts with the lipid bilayer.^[18–20]^ The presence of a lipid bilayer has been shown to influence these cytP450-redox partner protein-protein interactions both through dictating favorable orientations for complex formation, increasing or decreasing electron transfer rates and affecting metabolism of substrates, and altering the spin state shifts of the heme group of cytP450 when the redox partner is present.^[2, 21, 22]^ In order to study this ternary complex in a lipid environment, herein we utilized nanodiscs (ND). Using 4F peptides as the scaffold belt for the NDs is very favorable for this ternary complex because it creates very flexible nanodiscs that are accommodating for the stepwise addition of single-pass transmembrane proteins. Our previous studies have illustrated this ability with two membrane proteins.^[10]^

**Figure 1.**
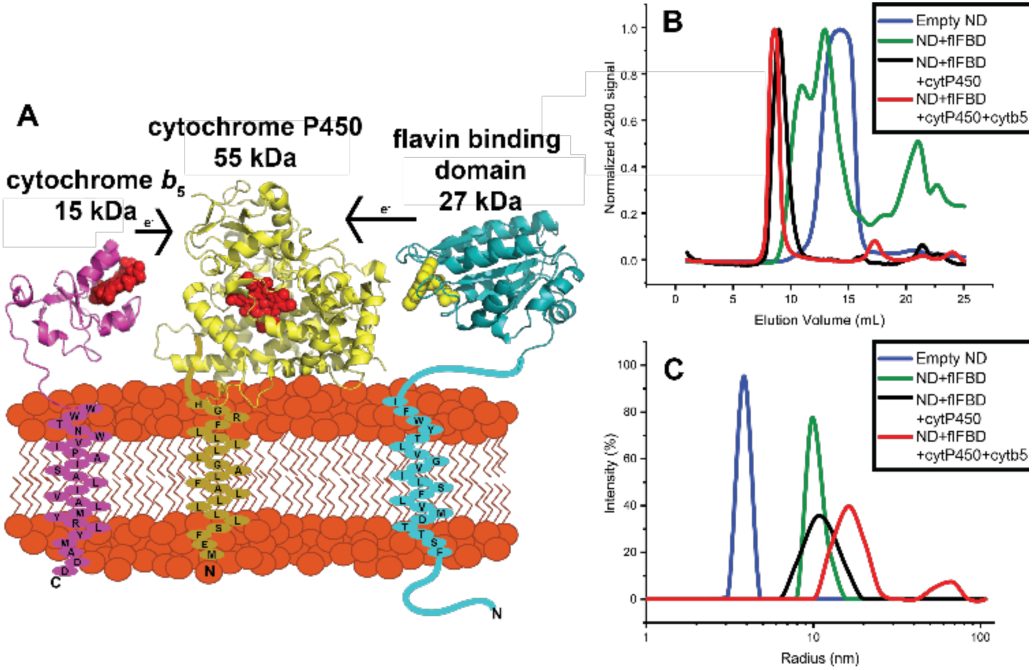
Reconstitution of cytochrome proteins in a lipid nanodisc. (A) A schematic of the three cytochrome proteins incorporated into a nanodisc. (B) and (C) both display the formation of protein-protein complexes into nanodiscs with stepwise incorporation of proteins into the nanodisc through SEC (B) and DLS (C).

As substrates have been shown to have a sizeable impact on strengthening the complex interactions between cytb_5_ and cytP450, we chose five substrates to probe this complex’s strength. These substrates were chosen due to their range in hydrophobicity although as cytP450 substrates, they are all hydrophobic in nature, and availability of cytP450 crystal structures solved in the presence of these compounds. They are: butylated hydroxytoluene (BHT; LogP = 5.3), bifonazole (BFZ; LogP = 4.8), benzphetamine (BZ; LogP = 4.1), 4-(4-Chlorophenyl)-1H-imidazole (4-CPI; LogP = 2.4), and 1-(4-Chlorophenyl)-imidazole (1-CPI; LogP = 2.3). Herein, we also examined the role of membrane and various substrates on the interplay between cytP450 and its two redox partners.

In order to study the ternary complex in lipid NDs, each full-length protein was expressed, purified and characterized as detailed in the SI. For this study, the 55 kDa cytP450 isoform that was utilized is cytP450 2B4, a rabbit homolog with 76% sequence identity to human cytP450 2B6.^[23]^ Sequential incorporation of the three proteins was accomplished over the course of three days using 4F-DMPC NDs that were purified through size exclusion chromatography (SEC). Two versions of the ternary complex were assembled: one with cytb_5_ incorporated into the ND first, then cytP450, and then flFBD, called “*cytb*_5_-*ternary complex*” and the other with flFBD incorporated into the ND first, then cytP450, and then cytb_5_, called “*flFBD-ternary complex*”. Reconstitution of these proteins into NDs was accomplished by the mixing of empty NDs and protein, incubating overnight, purifying by SEC, and characterizing their size by dynamic light scattering (DLS). On the second day, one molar equivalent of cytP450 2B4 was added to form complexes of cytb_5_-cytP450 or flFBD-cytP450 in NDs. On the third day, the other redox partner was added to the samples in order to make the *cytb_5_-ternary complex* or *flFBD-ternary complex* for a final protein ratio of 1:1:1. In Figure 1B and 1C, a gradual increase in size is displayed while creating the *flFBD-ternary complex* as seen by the increase in the Stoke radius in DLS or elution time growing shorter in SEC after the addition of proteins to empty NDs (blue), NDs+flFBD (green), NDs+flFBD+cytP450 (black), and NDs+flFBD+cytP450+cytb_5_ (red). Figure S1A and S1B show the *cytb_5_-ternary complex* formation via SEC and DLS. Previous work has revealed that substrates drive the formation of a strong complex between cytb_5_ and cytP450 in solution as well as in ternary complex systems with cytb_5_, cytP450, and FBD.^[24–26]^ Complex formation between ^15^N-labeled cytb_5_ and cytP450 was monitored through ^15^N/^1^H TROSY HSQC NMR experiments by measuring the signal intensities and linewidths of cytb_5_’s amide ^15^N-1H peaks. As the chemical environment around amide N-H changes upon binding to cytP450, a sizeable change in linewidth and intensity of resonances were observed in the NMR spectra (Figures S2-S14).

Figure 2 reports the average signal intensity observed for residues clustered on the lower cleft of cytb_5_. These residues (N62-R73) are highlighted in blue in the inset of Figure 2 and have been identified as important residues that bind to cytP450.^[27]^ As this lower cleft is highly involved in binding to cytP450, it is a good marker to probe if cytb_5_ is bound to cytP450. Upon the addition of cytP450 (Figure 2, orange) to the ^15^N-labeled cytb_5_, the overall signal intensity of the lower cleft residues of cytb_5_ drops to about ~52% of the original intensity. Intriguingly, when various substrates were added to the cytb5-cytP450 complex, the binding is not strengthened based on the lack of changes observed for the signal intensities (Figure 2, Figure S15). This lack of substrate effect is not completely surprising because the complex is already showing a tight binding due to the presence of the lipid membrane in the nanodiscs in comparison to the lipid-free condition as reported previously^[24]^. One hypothesis as to why the substrates do not increase the complex strength is that the hydrophobic compounds do partition into the lipid bilayer. As the drugs have another way of increasing hydrophobic contacts than promoting binding between the two proteins, the substrate-induced changes in the protein-protein interaction does not happen as strongly as observed in the absence of lipids. Some drugs like BFZ and BZ, slightly dislodge the complex between cytb_5_ and cytP450 (Figure 2 light green, light blue) as shown by the slight increase in signal intensity. While we do not know what the cause for this disruption is, there could be several things happening. As hydrophobic compounds partition into the membrane, they could be interacting with the transmembrane domains of the proteins or with other parts of cytP450.

**Figure 2:**
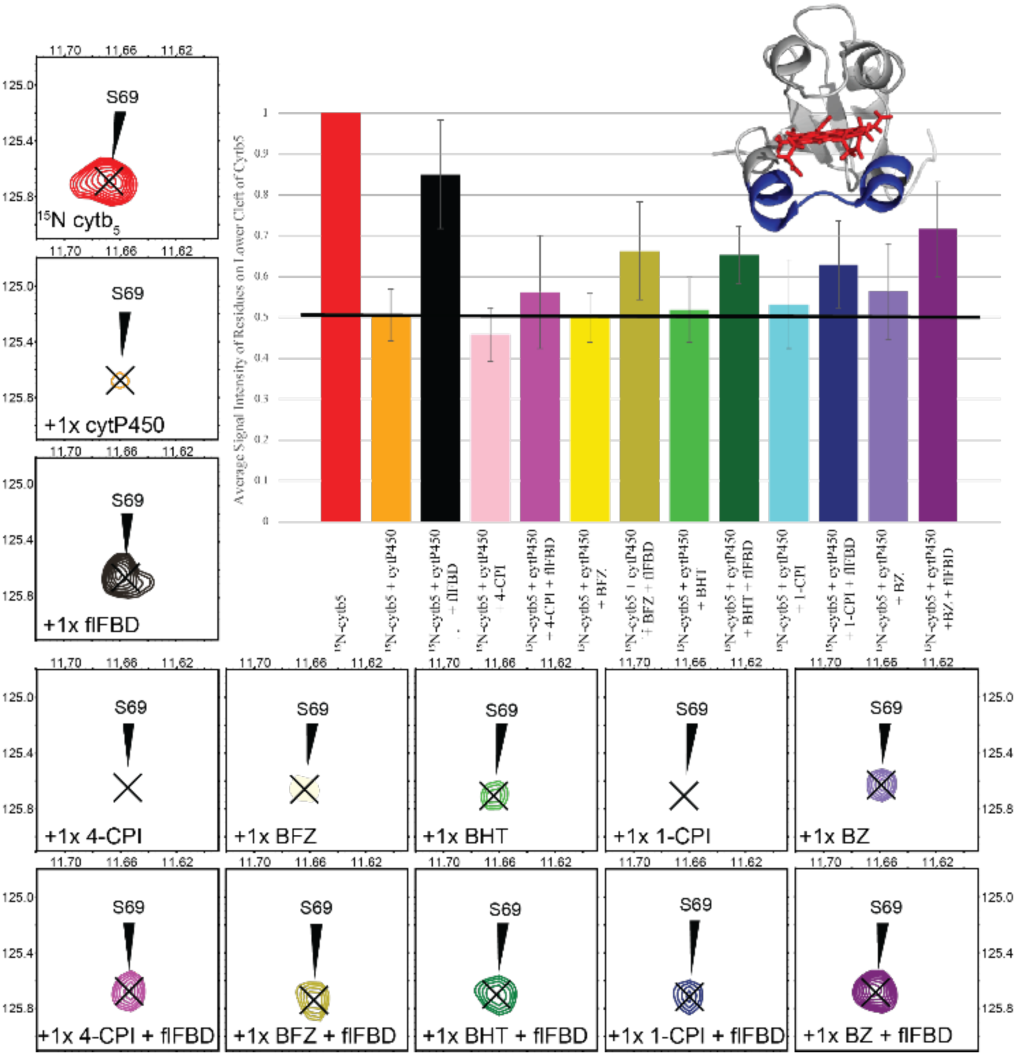
^15^N-cytb_5_ monitored ternary complex formation in nanodiscs. Average signal intensity of cytb_5_ residues involved in binding to cytP450 (N62-R73) measured from TROSY-HSQC spectra.^[28]^ Each bar in the graph as well as the contour peak in a 2D TROSY-HSQC spectrum corresponds to the addition of a protein or drug to the NDs containing ^15^N-labeled cytb_5_ sample. Peak intensity observed from nanodiscs containing ^15^N-labeled cytb_5_ alone was used as the reference (red) and set to 100%. The peak intensities were greatly reduced when cytP450 was added to the cytb_5_ in ND (orange). After flFBD was incorporated into the ND containing cytb_5_ and cytP450 (black), signal intensities were partially restored. Signal intensity depicting the effect of each of the five drugs is represented in the lighter shade bar and the darker shade bars are the measurement after the addition of flFBD to the substrate bound-cytb_5_-cytP450 complex: 4-CPI (pink); BFZ (yellow); BHT (green); 1-CPI (blue); BZ (purple). The average signal intensity represented by the black horizontal line, (inset) The lower cleft residues are highlighted in blue on a structure of cytb_5_ (2M33). Error bars were generated from the standard deviation of the average signal intensities of the mentioned residues.

Upon the addition of flFBD to the ND containing cytb_5_ and cytP450, we see an increase in signal intensity globally, but the cytb_5_ residues do not return to the same intensity level as observed for NDs containing cytb_5_ alone. In the presence of substrates, flFBD is unable to greatly disrupt the interaction between cytb_5_ and cytP450. Without a substrate present, flFBD can dislodge cytb_5_ from a complex with cytP450 which is demonstrated by the return of about 85% of the starting signal intensity. By dislodging cytb_5_, we mean that flFBD is able to disrupt the complex between cytb_5_ and cytP450 and interfere so some of the cytb_5_ is no longer bound to cytP450. Cytb_5_ does not leave the ND. Substrates can keep cytb_5_ and cytP450 bound to one another and make flFBD less capable of disrupting the complex. Drugs with highest to lowest ability of maintaining complex formation are in the following order: BZ, BFZ, BHT, 1-CPI, 4-CPI. Looking closely at one of the residues identified in binding of cytb_5_ to cytP450, Serine 69, we can see in Fig. 2 that its signal intensity decreases upon the addition of cytP450. After the addition of drugs to the cytb_5_-cytP450 ND complex, slight changes occur in the signal intensity. Once flFBD was added, varying levels of signal intensities were restored. Figure S15 displays the extracted linewidths of S69 to illustrate the broadening of the signal upon the addition of cytP450 and the partial restoration after the addition of flFBD.

From our previous study ^[24]^, it was shown that cytb_5_ can dislodge FBD from the FBD-cytP450 complex. We were curious as to how the membrane would affect cytb_5_’s ability to disrupt the FBD-cytP450 complex as it provides both a more native membrane environment and a spatial constraint (Figure 3). Uniformly ^15^N-labeled flFBD was expressed, purified, and reconstituted into 4F-DMPC NDs. A ^15^N/^1^H TROSY-HSQC spectrum was acquired of ^15^N-flFBD in ND alone (red). Stepwise reconstitution was done to incorporate cytP450 2B4 into the flFBD containing NDs and a spectrum was obtained (orange). Each of the five drugs chosen were incubated with the complex of flFBD-cytP450 (4-CPI, pink; BFZ, yellow; BHT, green; 1-CPI, blue; BZ, purple) and then TROSY-HSQC spectra were obtained (Figures S16-S28). Finally, cytb_5_ was incorporated into the flFBD-cytP450-ND with and without substrates.

**Figure 3:**
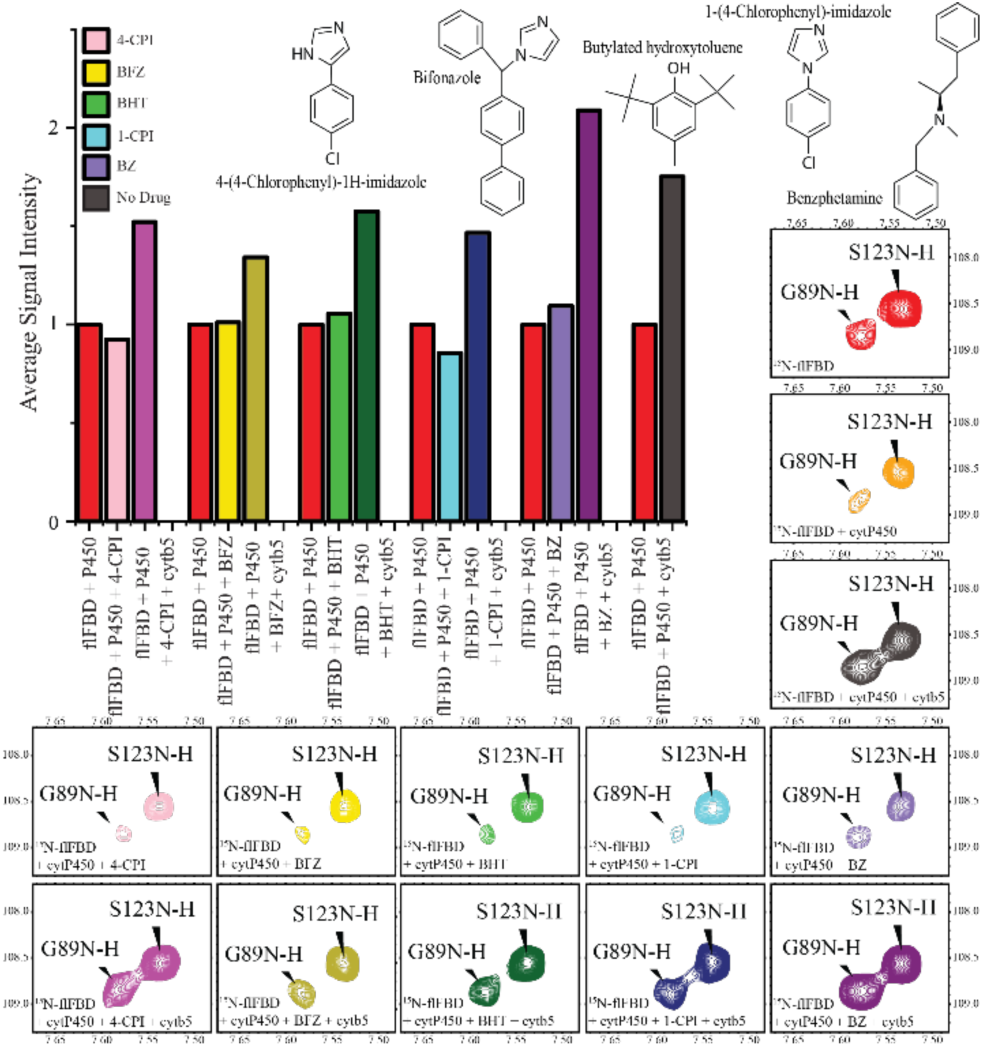
^15^N-flFBD monitored ternary complex formation demonstrates cytb_5_’s ability to dislodge flFBD from its complex with cytP450. The bar graph shows the average overall signal intensity of all ^15^N-labeled flFBD residues. In red is the complex intensity at 100% for flFBD in complex with cytP450. After the red bar is the intensity for the complex with a substrate added (4-CPI, BFZ, BHT, 1-CPI, or BZ) or no substrate. The third bar is the addition of cytb_5_ to the complex with a substrate. Highlighted sections of ^1^H/^15^N TROSY-HSQC spectra of ^15^N-flFBD displaying residue G89 which has been shown to be involved in binding to cytP450 and S123 which is not involved, (red) ^15^N-flFBD; (orange) ^15^N-flFBD with lxcytP450 2B4; (gray) ^15^N-flFBD with lxcytP450 2B4 and lxcytb_5_. The chemical structure of each of the five substrates used is displayed in the middle with the next column showing the ^15^N~flFBD~cytP450 complex with substrate spectrum and the rightmost column showing the ^15^N-flFBD-cytP450-substrate-drug spectrum. 4-CPI (pink); BFZ (yellow); BHT (green); 1-CPI (blue); BZ (purple).

The overall signal intensity of the flFBD residues was monitored as the protein complexes were formed and broken. A decrease in the signal intensity upon the addition of cytP450 indicates that a complex was formed between flFBD and cytP450. After the addition of a substrate, no significant difference in complex strength was observed. Substrates have not been shown to dramatically increase the affinity of cytP450 for FBD, so this result was not surprising. Both 4-CPI and BFZ marginally increased the complex strength while BHT and BZ’s addition did not strengthen the complex, and BZ weakened it a little. The addition of 1-CPI, however, significantly increased the complex formation which is shown in Figure 3 (blue) and Figure S33.

Once cytb_5_ was incorporated into NDs, it does show the ability to disrupt the flFBD-cytP450 complex. This is demonstrated by the restoration of signal intensities of flFBD residues, particularly, looking at Glycine 89 (in Figure 3) which is a residue in loop 1 of flFBD which coordinates the FMN cofactor and has been previously implicated in binding to cytP450.^[10]^ By analyzing the signal intensity and linewidth of this residue, we obtained information about its chemical environment and the timescale of its dynamics. Upon addition of cytP450, the resonance broadens, and the intensity decreases as can be seen in Figure 3 (orange). Adding in a substrate did not significantly change the peak intensity for any of the substrates chosen for this residue of flFBD. Upon cytb_5_ incorporation into the nanodisc, G89’s linewidth and signal intensity were greatly restored (bottom row, Figure 3). Meanwhile other residues on flFBD that are farther away from the interaction like S123, undergo some changes but these are much more consistent with the nanodisc containing a larger complex’s tumbling rate rather than direct interaction with cytP450. Figure 4 displays the extracted linewidths of G89 and other important loop residues to illustrate the broadening of the signal upon the addition of cytP450 to the flFBD in ND and the restoration after the addition of cytb_5_ to the complex in NDs.

**Figure 4.**
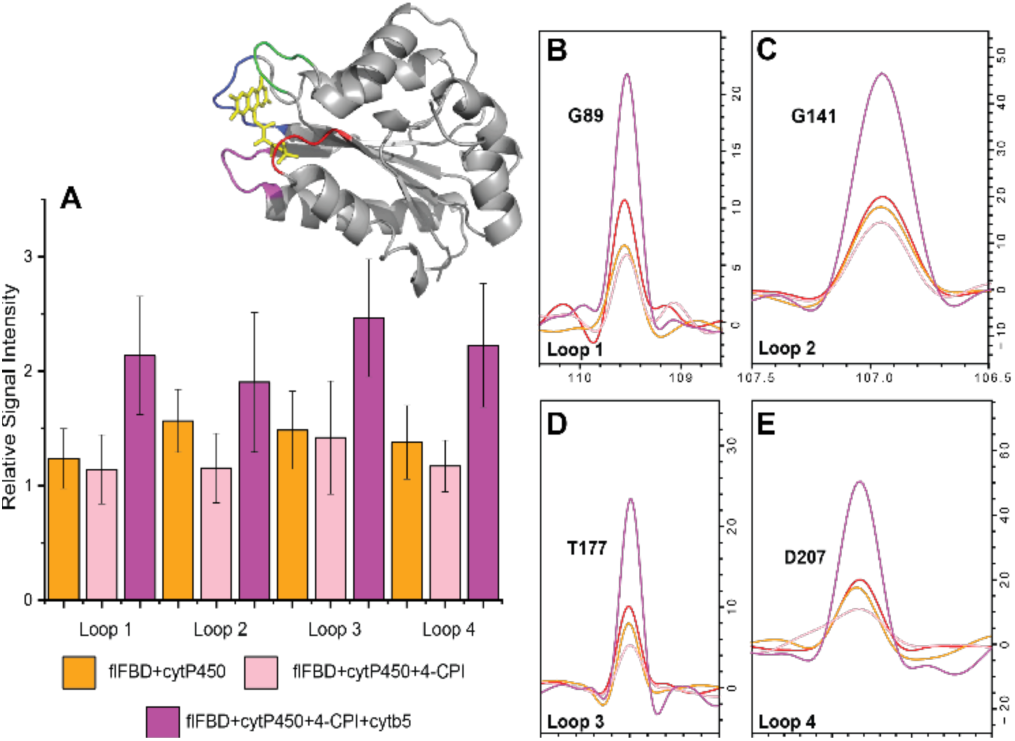
Loops of flFBD reveal changes in its complexation state. (A) Changes in signal intensities measured from TROSY-HSQC spectra of flFBD for its four loop regions which coordinate its FMN cofactor. Each color represents the different states of flFBD as indicated. (inset) The loop regions in the structure are highlighted with loop 1 (red), loop 2 (green), loop 3 (blue), and loop 4 (magenta). B-E. ^15^N spectral slices extracted from 2D TROSY-FISQC spectra reveal broadening and restoration of signals for representative peaks of each loop; red trace is for flFBD alone in the ND. The error bars were determined from the standard deviation of the average from the selected residues of each loop (Table S1).

Studying the ternary complexes incorporated into lipid nanodiscs allowed us to capture some interesting behavior of cytP450’s preference for its redox partners. While cytb_5_ is still able to dislodge flFBD from a complex with cytP450, it is not to as great of an extent as it could in the absence of lipids in solution. Having the three proteins being spatially confined in a nanodisc could contribute to keeping flFBD bound to cytP450 better, but it is currently unclear to assign that to the influence of lipids on complex strength or the physical restraint of motion. The variety of substrates utilized in this study all behaved similarly in regard to forming a stronger complex with cytb_5_-cytP450, however, it was much less of an effect than what was seen in lipid-free solution ^[23]^. flFBD was able to dislodge cytb_5_ from its complex with cytP450, but not completely. The highest effect was seen without substrates and flFBD was only able to slightly disrupt the complex in the presence of substrates.

One of the more interesting comparisons from this study includes the two inhibitors 4-CPI and 1-CPI in the presence of ^15^N-*flFBD-ternary complex*. In agreement with Zhao et al., we have found major differences in the interactions between cytP450 and flFBD in the presence of 1-CPI versus 4-CPI, even with their structural similarity.^[28]^ Crystal structures of cytP450 2B4 have been solved in the presence of both inhibitors which revealed very different active site conformations. The calculated active site volume for 4-CPI was 200 Å versus 280 Å for 1-CPI bound cytP450 2B4. Our NMR data reveal that there is an increased binding and tighter complex formation between cytP450 2B4 and flFBD after the 1-CPI inhibitor addition which is in agreement with the ITC experiments performed by Zhao et al.^[28]^ Figure 4 and Figures S29-S31 show the average signal intensities for each of the four loops that coordinate the FMN cofactor. The residues in each loop are listed in Supplemental Table 1. In the presence of 4-CPI binding to cytP450, all loops are only slightly affected. In comparison, in the presence of 1-CPI binding to cytP450, loop 2 on flFBD is strongly affected. Even the addition of cytb5 does not completely restore the signal intensity of loop 2 in the 1-CPI complex sample.

Interestingly this data reveals more residues binding of flFBD to cytP450 which was recently reported.^[9]^ In the presence of membrane, flFBD binds to cytP450 with all four loops that surround the FMN. The major takeaway from this study is that in the presence of lipid membrane, this dynamic interplay approaches what is thought to be more physiological behavior. Without being reconstituted in a lipid nanodisc, flFBD cannot dislodge cytb_5_ from binding to cytP450. Physiologically, if cytb_5_ is bound to cytP450 before the first electron can be donated by CPR, then the catalytic cycle ceases. Cytb_5_ is incapable of donating the first electron due to the disparity in redox potentials. Without the dynamic exchange between redox partner binding, drug metabolism slows or even halts. Here we see signs that add to the prevailing hypothesis that cytb_5_ plays not only an electron donating role, but also has conformational contributions to influencing cytP450’s behavior. A myriad of studies has illustrated that while holo-cytb_5_ has the largest effect, mainly an increase, in substrate turnover, apo-cytb_5_ as well as other metal substituted porphyrins, also have a sizeable contribution to cytP450’s substrate metabolism that is independent of the electron transfer ability.^[29–31]^ In this study, we carry out all NMR experiments with oxidized proteins and an absence of reductants. Due to the inability of cytb_5_ to donate electrons to cytP450, we can assume that some conformational effects are at play.

In conclusion, as we have shown through NMR results, the dynamic exchange between cytP450, flFBD, and cytb_5_ is highly affected by the presence of a lipid environment. We see cytb_5_ dominating the competition by weakening the flFBD’s complex with cytP450. In the membrane environment, flFBD is more able to disrupt the cytb_5_-cytP450 complex than in the absence of lipids albeit not completely. The addition of substrates to this ternary complex reveals differences in redox partner binding to cytP450 as well as the skewing of the interplay to favor cytb_5_ over flFBD binding. This study emphasizes the need to study these proteins in a membrane environment in order to more fully capture their native behaviors.

## Acknowledgements

This study was supported by NIH funding (GM084018 to A.R.).

