## Supplemental figures and table for "Probing protein-protein and protein-substrate interactions in the dynamic membrane-associated ternary complex of Cytochromes P450, b5, and Reductase"

### Materials and Methods

#### *Materials and Reagents*

*E. coli* C41 cells for protein overexpression were purchased from Lucigen (Middleton, MI). Yeast extract, tryptone, and sodium chloride for unlabeled growth media were purchased from Sigma-Aldrich. $^{15}N$ Ammonium Chloride and $D_2O$ were purchased from Cambridge Isotope Laboratories (Andover, MA). I,2-dimyristoyl-sn-glycero-3-phosphocholine (DMPC) lipid was purchased from Avanti Polar Lipids (Alabaster, AL). 5-mm symmetrical $D_2O$-matched Shigemi NMR microtubes were purchased from Shigemi, Inc. (Allison Park, PA). Resins, phosphate buffer components (monobasic and dibasic), and all other chemicals were purchased from Sigma-Aldrich.

#### *Expression and Purification of full-length cytochrome b5*

Full-length uniformly $^{15}N$-labeled and unlabeled wild-type rabbit cytb5 was expressed and purified as described previously. [1-3] Briefly, *E. coli* C41 cells were transformed with a pLW01 plasmid containing the cytb5 gene. The cells were grown up in LB medium to an OD of 1 (at 600 nm). This culture was diluted 100-fold into 100 mL of $^{15}N$-Celtone medium. Then the culture was grown at 35 °C under shaking at 250 rpm until an OD of 1 at 600 nm was achieved. The cells were pelleted and resuspended in 10 mL of fresh $^{15}N$-Celtone medium. The resuspended cell culture was added to the final 1 L of culture minimum medium. Isopropyl β-D-thiogalactopyranoside was added to a final concentration of 10 µM, and incubation was continued for 20 h, at which time the cells were harvested.

#### *Expression and Purification of full-length FBD*

U-$^{15}N$-labeled and unlabeled flFBD were expressed and purified as described previously [4]. The fl-FBD gene is encoded on a pSC plasmid and is preceded by the OmpA signal peptide [5]. Briefly, fl-FBD was expressed in *E. coli* C41 cells in either LB medium to obtain unlabeled protein, or M9 medium to obtain U-$^{15}N$ labeled samples, supplemented with 5 nM FMN. Protein expression was induced at OD600 = 0.7 by adding 0.4 mM IPTG to the cultures for 16 h at 30 °C. After cell harvest at 6000 x g and 4 °C, the cells were lysed by 30 µg/ml lysozyme and protease inhibitor in Tris-Acetate buffer pH 7.4 for 30 mins at 4 °C, followed by sonication (with 1 s on and 1 s off pulses) for 5 mins. The membrane fraction was pelleted by ultracentrifugation at 105,000 x g and 4 °C for 45 mins, and further treated with 0.3% (v/v) Triton X-100 for 16 h at 4 °C. The solubilized membrane proteins were purified by DEAE anion exchange chromatography twice. For this purpose, the protein was loaded onto the column and eluted using a NaCl gradient ranging from 0.2 M to 0.5 M in Tris-Acetate pH 7.4 containing 1 µM FMN, 0.3% (v/v) sodium cholate.

#### *Expression and Purification of full-length cytP450 2B4*

Full-length CYP2B4 was expressed and purified as described in the literature [1].

#### *Preparation of nanodiscs*

DMPC powder was suspended into buffer A (40 mM potassium phosphate, pH 7.4) to make a stock solution at 20 mg/mL. The 4F peptide (DWFKAFYDKV AEKFKEAF) was dissolved in buffer A to make a stock solution at 10 mg/mL. The DMPC stock solution was vortexed and sonicated three times for 30 s each to create a suspension, and vortexed thoroughly immediately before use. The stock solution was mixed together at a peptide:lipid ratio of 1:1.5 % w/w and incubated at 37 °C overnight with slow agitation. The nanodiscs were purified by size exclusion chromatography (SEC). A Superdex 200 Increase 300/10 GL column was operated on an AKTA purifier (GE Healthcare, Freiburg, Germany).

#### *Reconstitution of full-length proteins in nanodiscs*

Cytb5 or flFBD was added to the empty nanodiscs at molar ratios of 1:1.2 (protein/nanodisc) and incubated for 16 hours at 25 °C with gentle agitation. The reconstituted nanodiscs were further purified by SEC. Fractions showing absorbance at 417 nm (cytb5) or 454 nm (FBD) were pooled and used for further analysis. In order to form a complex, cytP450 was added to purified cytb5 or flFBD containing nanodiscs at a molar ratio of 1:1. After overnight incubation for 16 hours at 25 °C with gentle agitation and another round of purification either cytb5 or flFBD was added to form all three protein-containing nanodiscs.

The empty nanodiscs and reconstituted proteins were subjected to dynamic light scattering (DLS) measurements on a DynaPro NanoStar instrument (Wyatt Technology Corp., Santa Barbara, USA) at 25 °C for 10 acquisitions of 5 s each. DLS and SEC measurements confirm the increase of the hydrodynamic radius after stepwise incubation with cytb5/flFBD, fl-CYP2B4 and fl-FBD/cytb5.

#### *NMR experiments*

NMR experiments were performed at 298 K on an 800 MHz Bruker spectrometer equipped with an Ascend magnet and TCI cryoprobe. 2D $^{1}H/^{15}N$ TROSY HSQC NMR spectra were recorded from 0.1 mM $^{15}N$-labeled protein (either flFBD or cytb5) in 40 mM potassium phosphate, pH 7.4. All NMR spectra were obtained using 128 scans and 128 t1 increments. Data was processed using TopSpin (Bruker) and analyzed with Sparky (Goddard). The previously reported cytb5 and flFBD backbone chemical shift assignments were used in this study.[6, 7]

### Supplemental Figures:

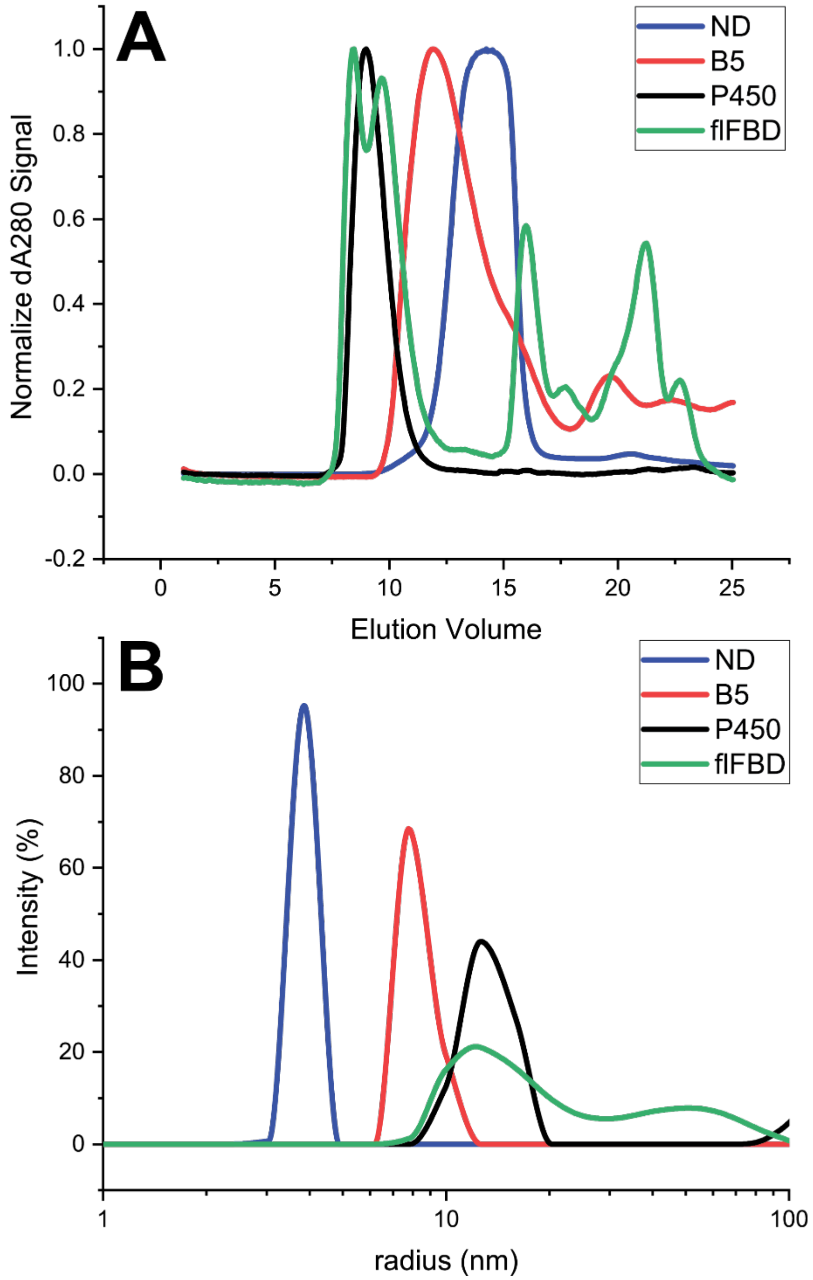


**Figure S1. Incorporation of the cytb5-cytP450-flFBD ternary complex into lipid nanodiscs.** A and B both display the formation of the *cytb5-ternary complex* into nanodiscs with stepwise incorporation of the three proteins through SEC (A) and DLS (B) for empty 4F-DMPC nanodiscs (blue), nanodiscs with cytb5 (red), nanodiscs with cytb5-cytP450 (black), and nanodiscs with cytb5-cytP450-flFBD (green).

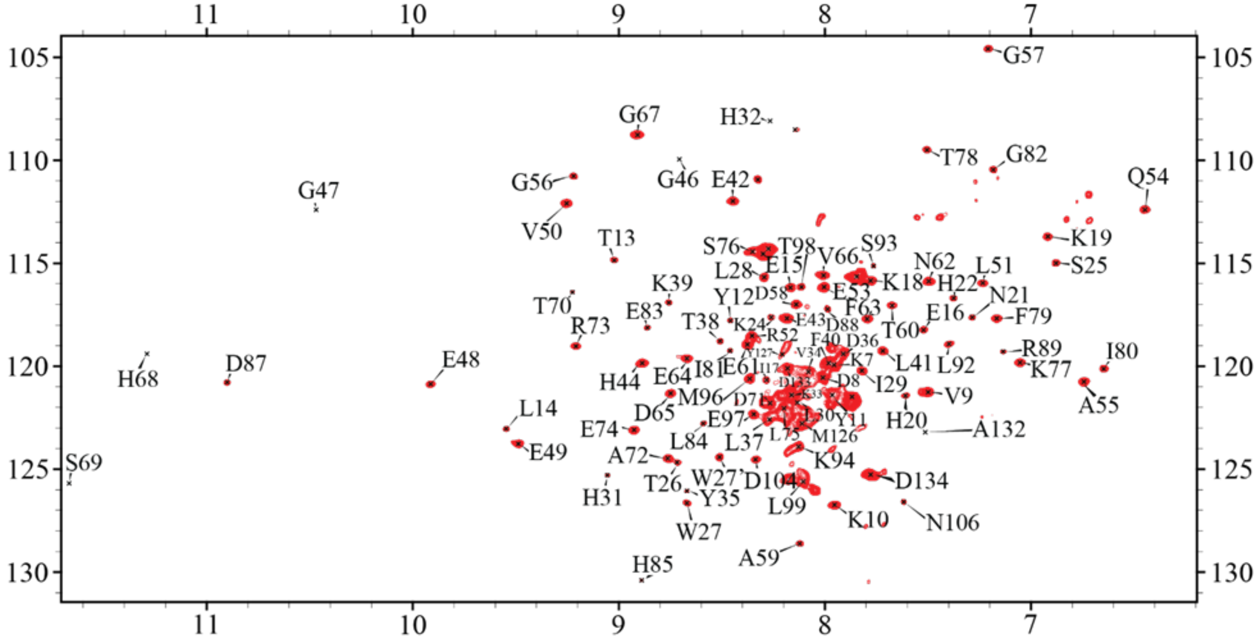


**Figure S2.** A 2D $^{1}H/^{15}N$ TROSY HSQC of $^{15}N$-cytb5 in 4F-DMPC ND.

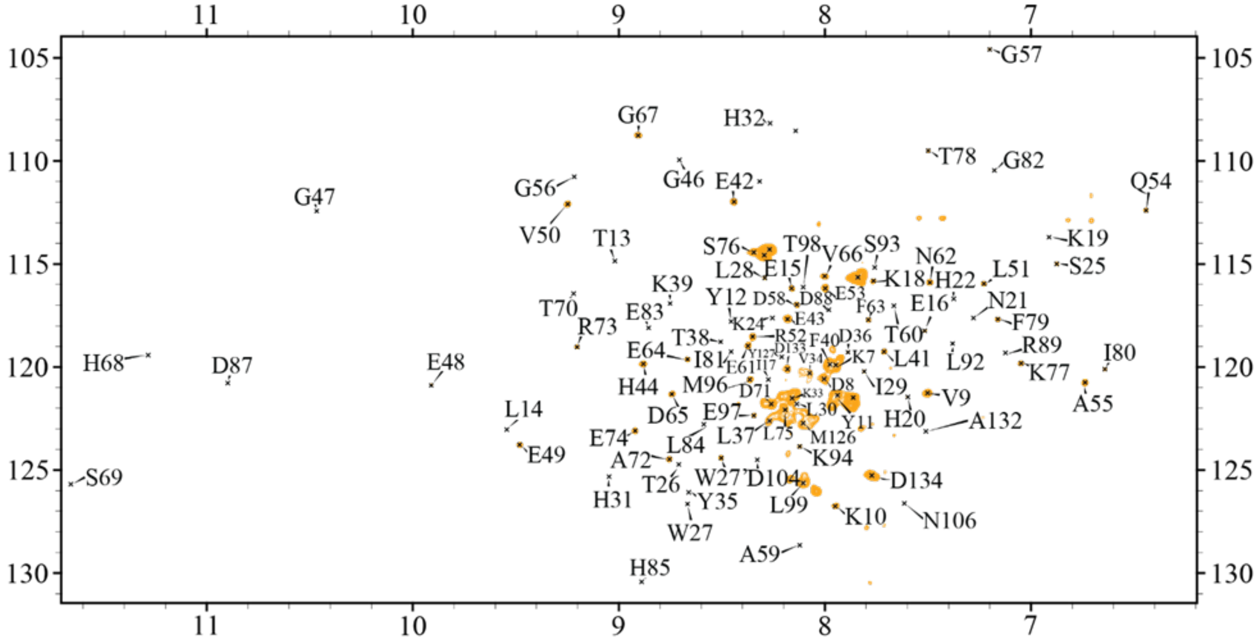


**Figure S3.** A 2D $^{1}H/^{15}N$ TROSY HSQC of $^{15}N$-cytb5 in 4F-DMPC ND with 1 molar equivalent of cytP450 2B4.

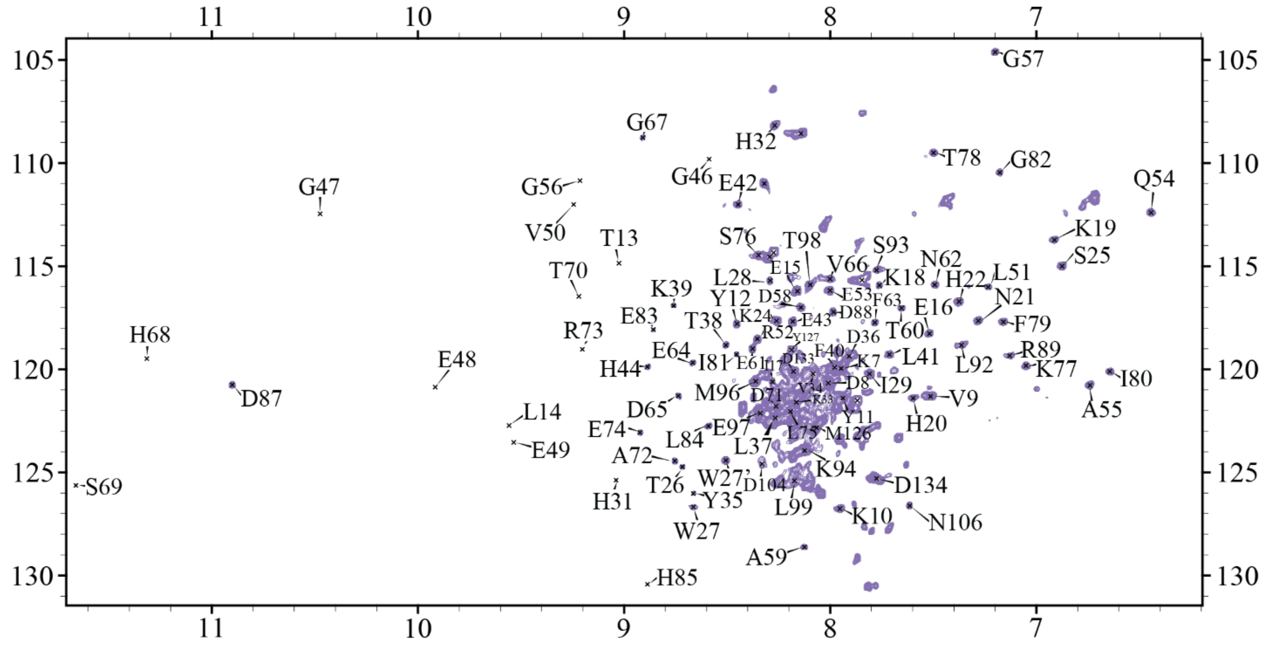

**Figure S4.** A 2D $^{1}H/^{15}N$ TROSY HSQC of $^{15}N$-cytb5 in 4F-DMPC ND with 1 molar equivalents of cytP450 and BZ.

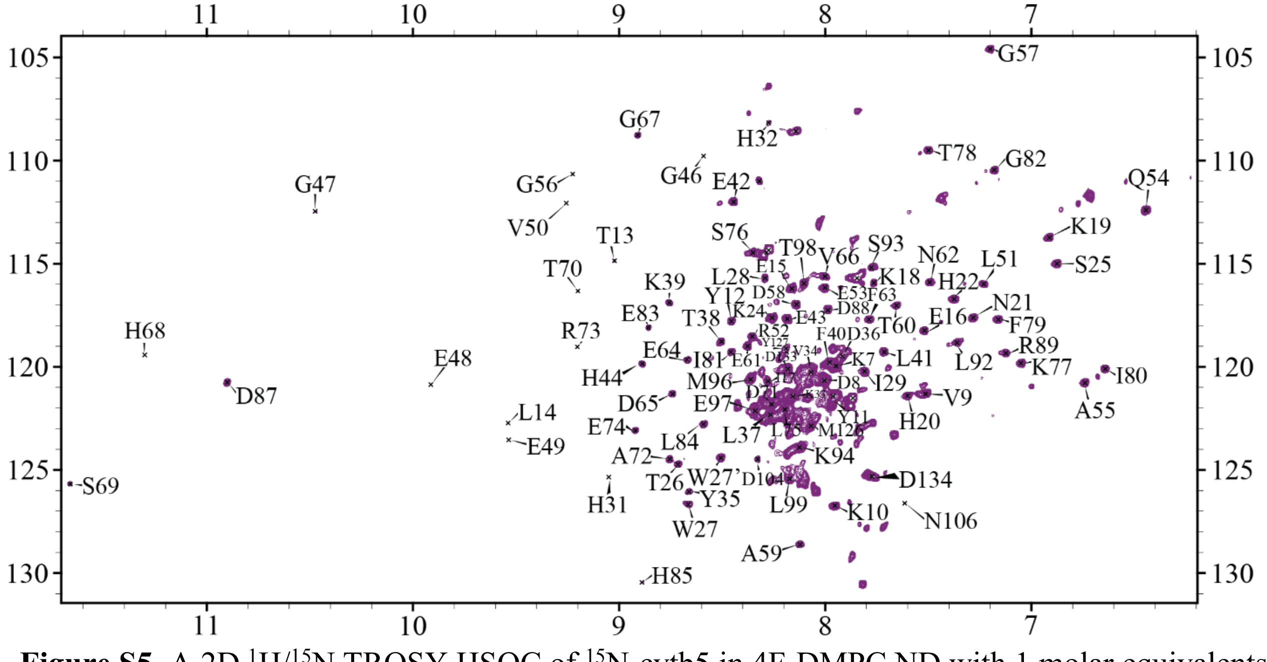

**Figure S5.** A 2D $^{1}H/^{15}N$ TROSY HSQC of $^{15}N$-cytb5 in 4F-DMPC ND with 1 molar equivalents of cytP450, BZ, and flFBD.

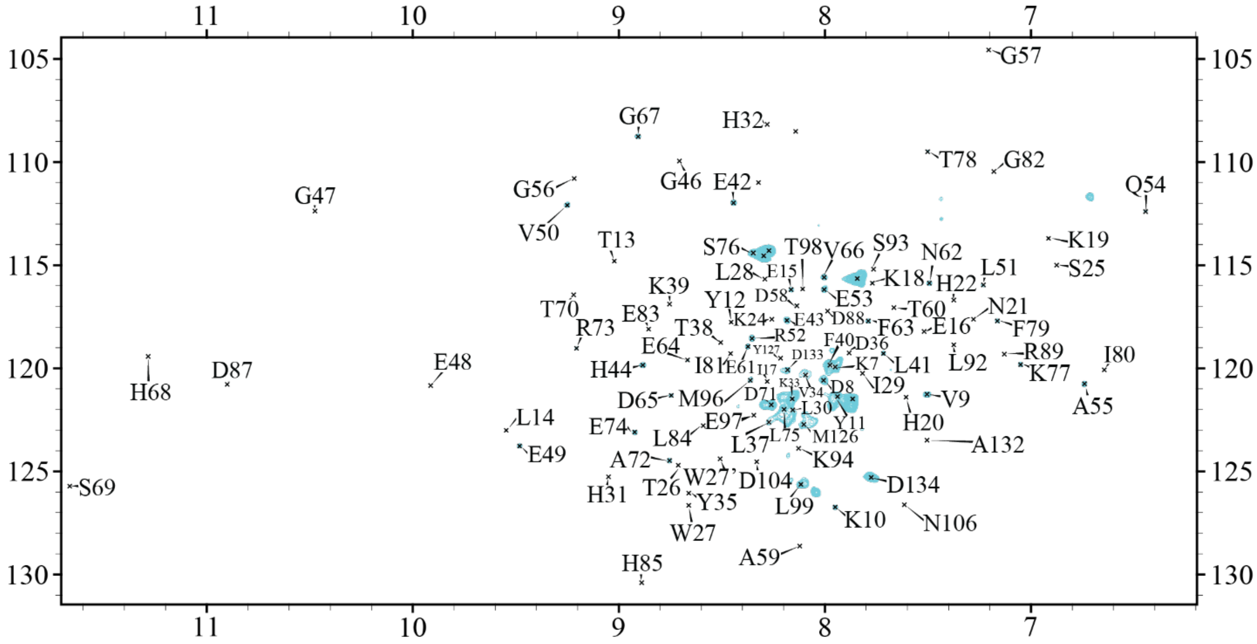


**Figure S6.** A 2D $^{1}H/^{15}N$ TROSY HSQC of $^{15}N$-cytb5 in 4F-DMPC ND with 1 molar equivalents of cytP450 and 1-CPI.

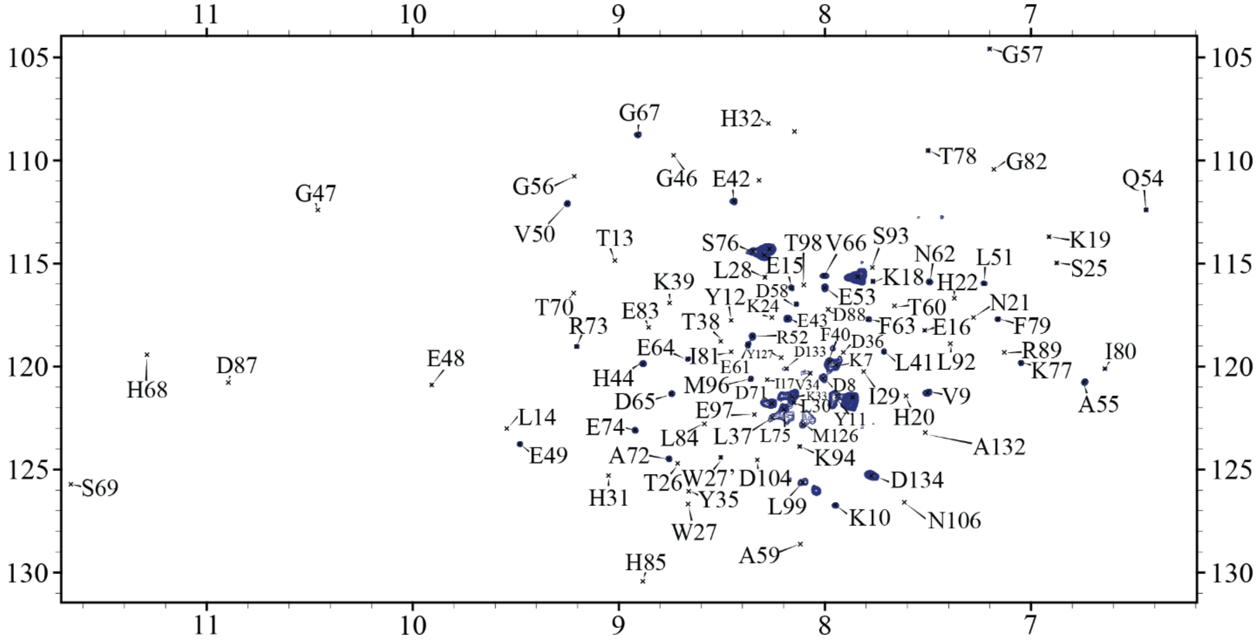


**Figure S7.** A 2D $^{1}H/^{15}N$ TROSY HSQC of $^{15}N$-cytb5 in 4F-DMPC ND with 1 molar equivalents of cytP450, 1-CPI, and flFBD.

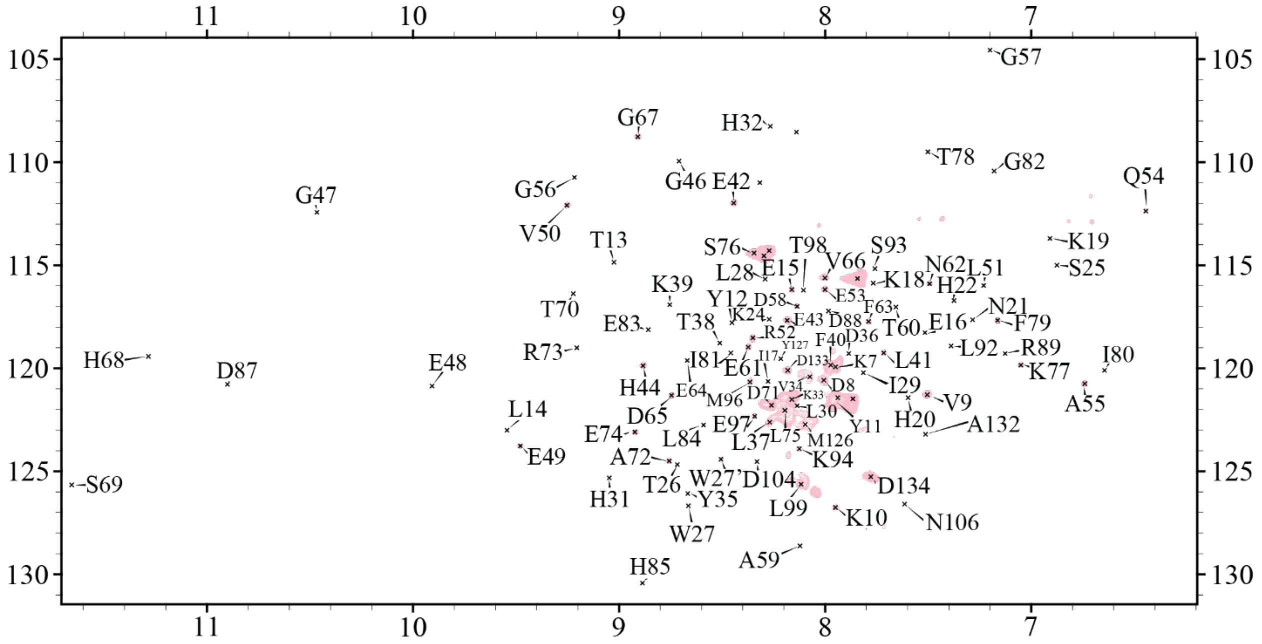


**Figure S8.** A 2D $^{1}H/^{15}N$ TROSY HSQC of $^{15}N$-cytb5 in 4F-DMPC ND with 1 molar equivalents of cytP450 and 4-CPI.

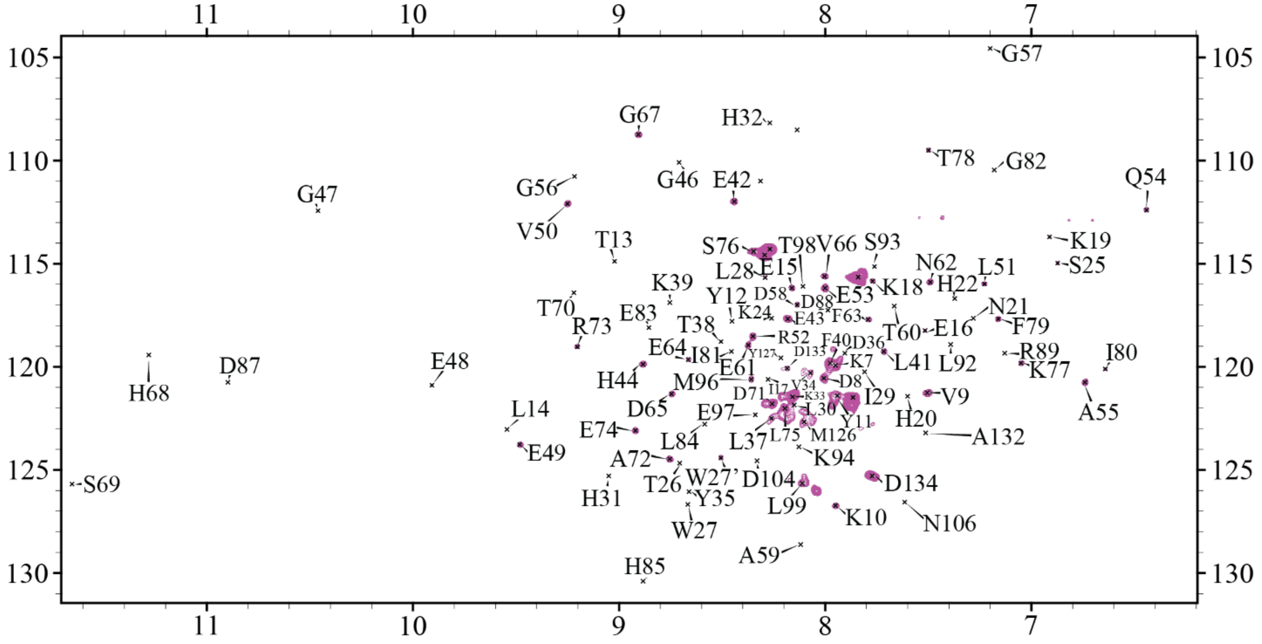


**Figure S9.** A 2D $^{1}H/^{15}N$ TROSY HSQC of $^{15}N$-cytb5 in 4F-DMPC ND with 1 molar equivalents of cytP450, 4-CPI, and flFBD.

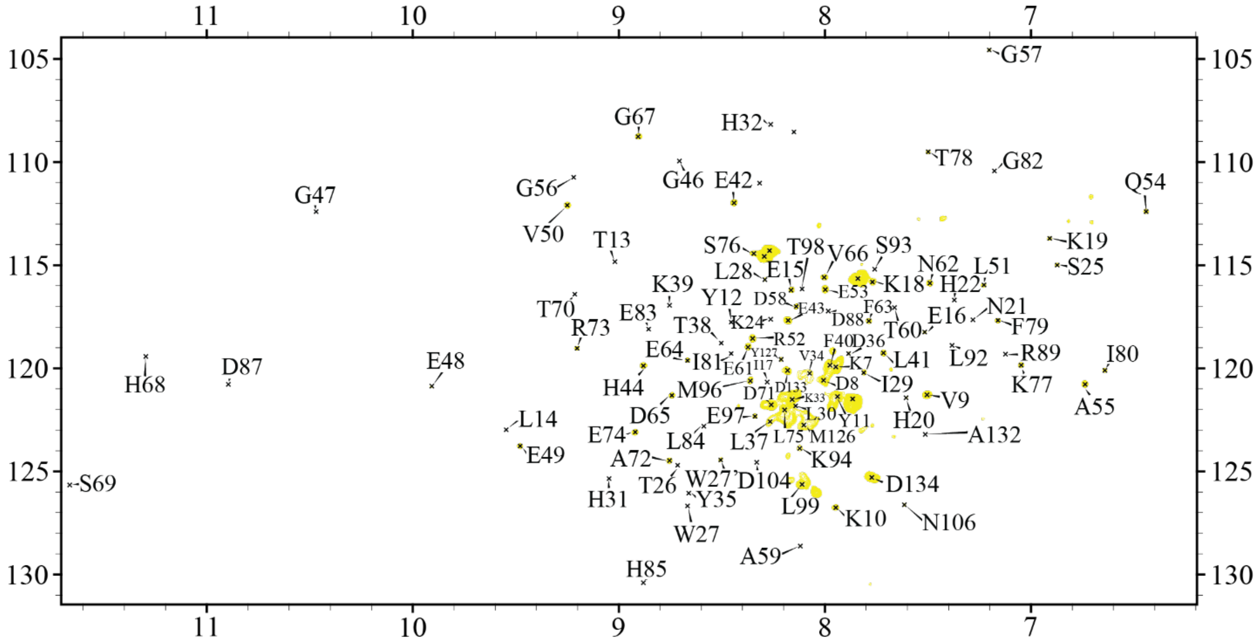


**Figure S10.** A 2D $^{1}H/^{15}N$ TROSY HSQC of $^{15}N$-cytb5 in 4F-DMPC ND with 1 molar equivalents of cytP450 and BFZ.

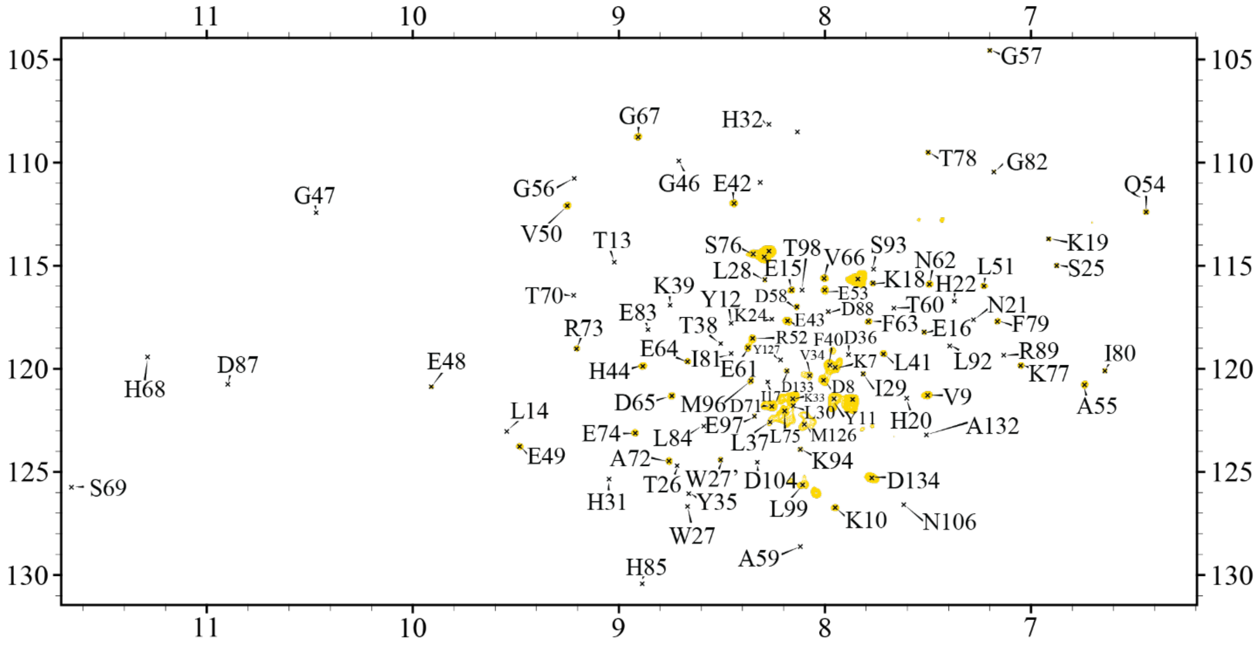


**Figure S11.** A 2D $^{1}H/^{15}N$ TROSY HSQC of $^{15}N$-cytb5 in 4F-DMPC ND with 1 molar equivalents of cytP450, BFZ, and flFBD.

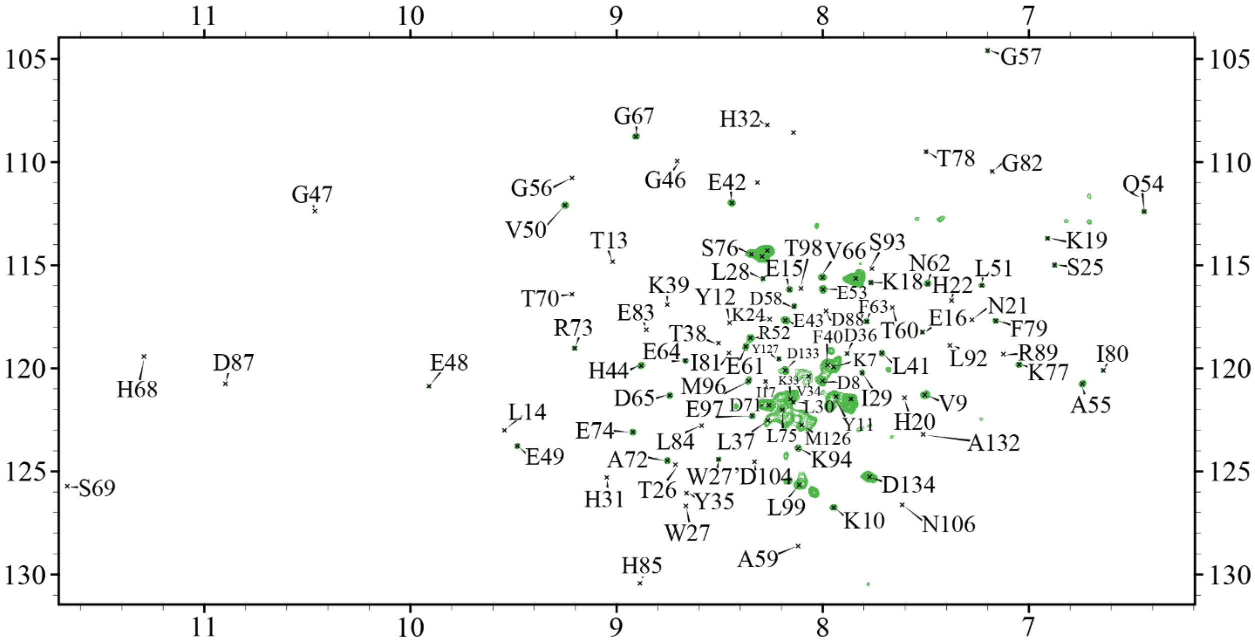


**Figure S12.** A 2D $^{1}H/^{15}N$ TROSY HSQC of $^{15}N$-cytb5 in 4F-DMPC ND with 1 molar equivalents of cytP450 and BHT.

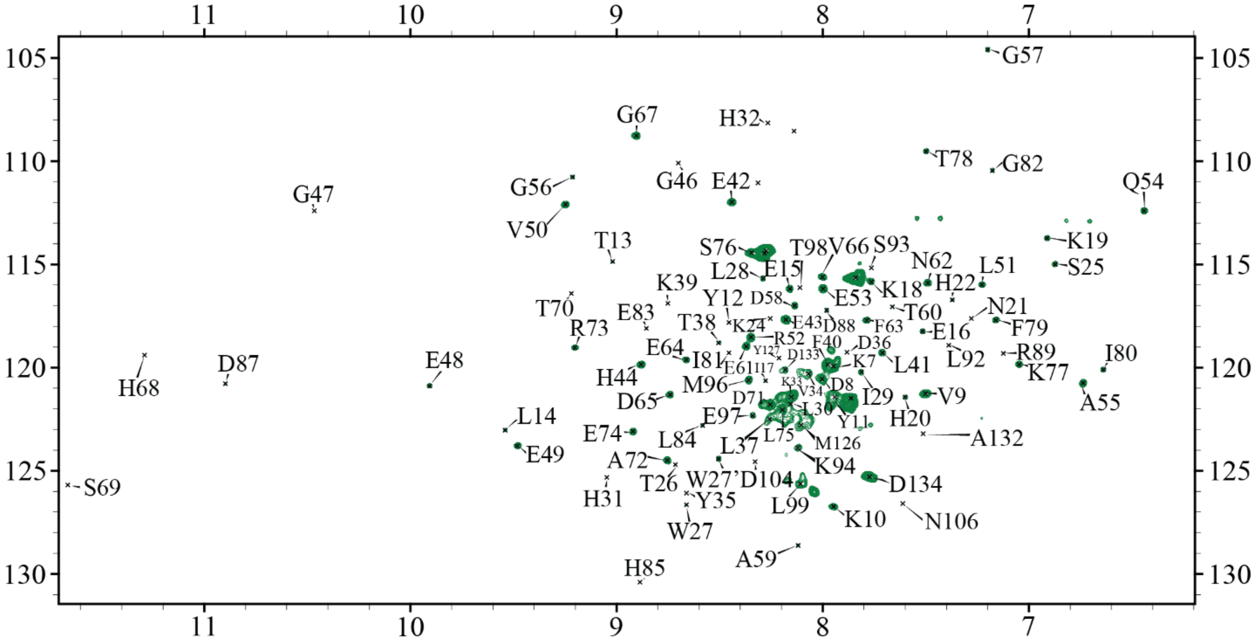


**Figure S13.** A 2D $^{1}H/^{15}N$ TROSY HSQC of $^{15}N$-cytb5 in 4F-DMPC ND with 1 molar equivalents of cytP450, BHT, and flFBD.

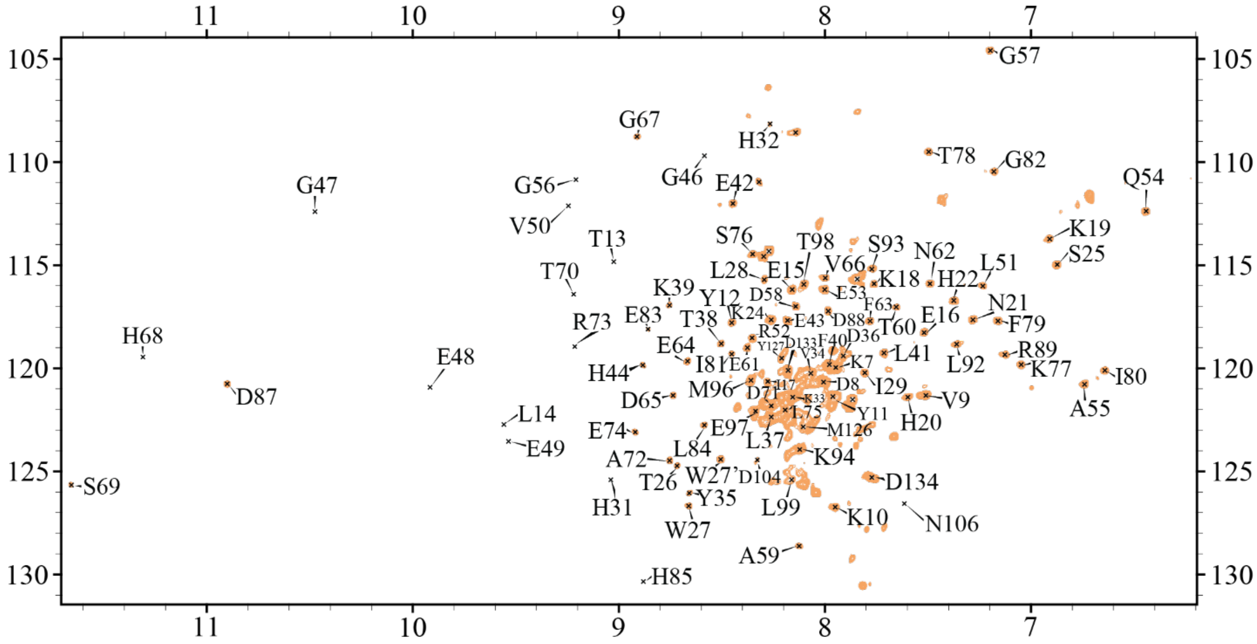


**Figure S14.** A 2D $^{1}H/^{15}N$ TROSY HSQC of $^{15}N$-cytb5 in 4F-DMPC ND with 1 molar equivalents of cytP450 and flFBD (no drug).

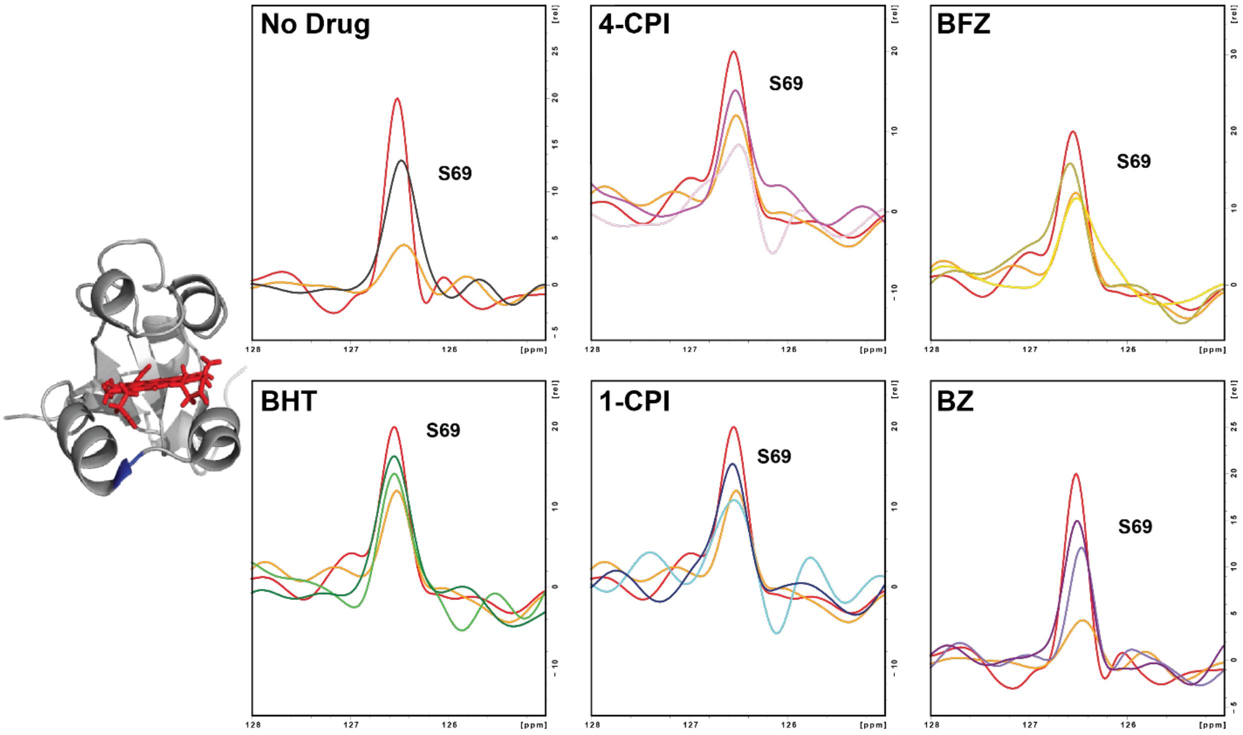


**Figure S15.** 1D line slices in the nitrogen dimension reveal line broadening and restoration of signal for a peak on cytb5's lower clef. As before, red is $^{15}N$-labeled cytb5 alone, orange is in complex with cytP450, light color is after the addition of drug, and dark color after the addition of flFBD for 4-CPI (pink), BFZ (yellow), BHT (green), 1-CPI (blue), and BZ (purple).

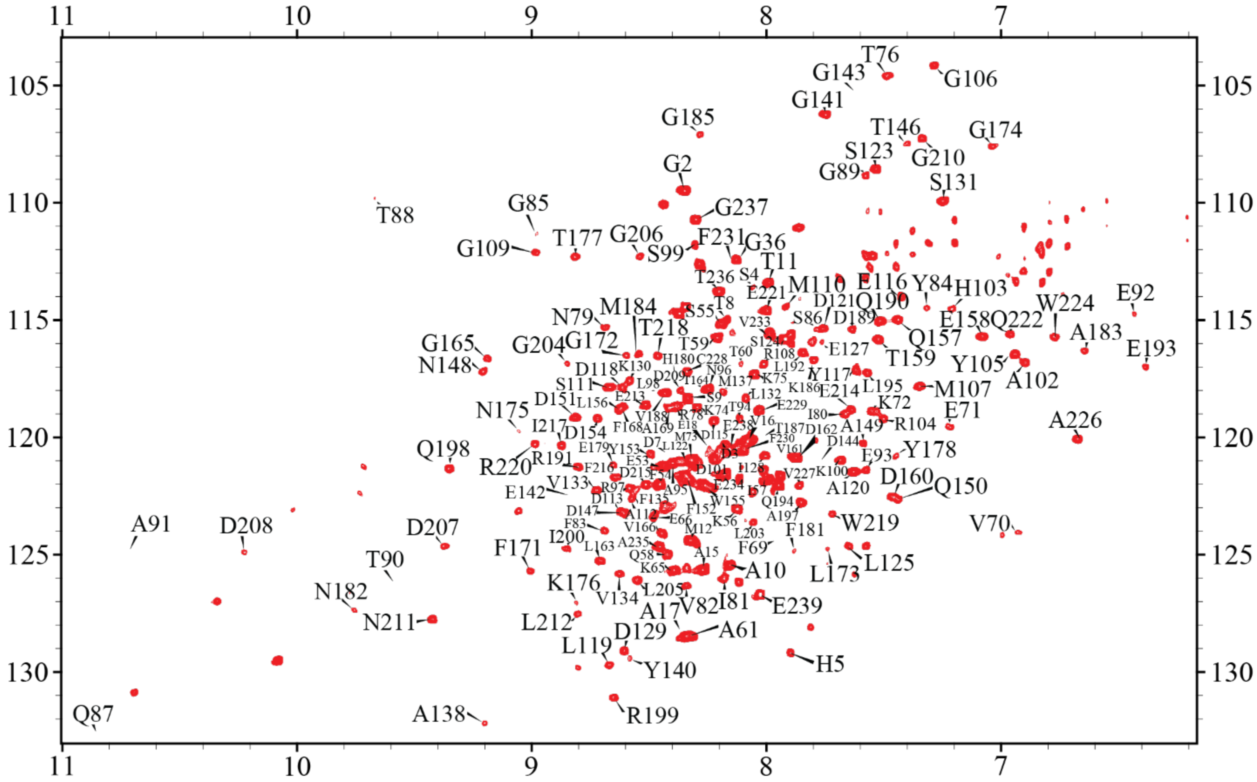


**Figure S16.** A 2D $^{1}H/^{15}N$ TROSY HSQC of $^{15}N$-flFBD in 4F-DMPC ND.

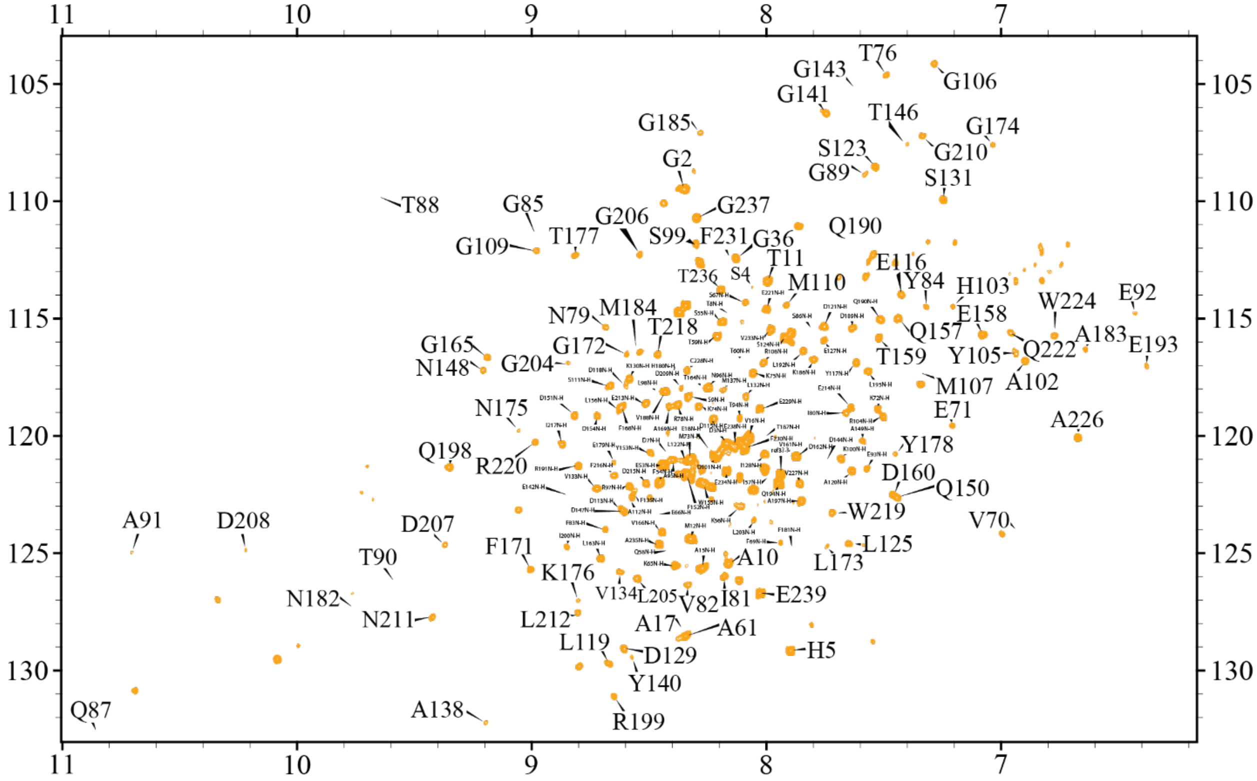


**Figure S17.** A 2D $^{1}H/^{15}N$ TROSY HSQC of $^{15}N$-flFBD in 4F-DMPC ND with 1 molar equivalent of cytP450.

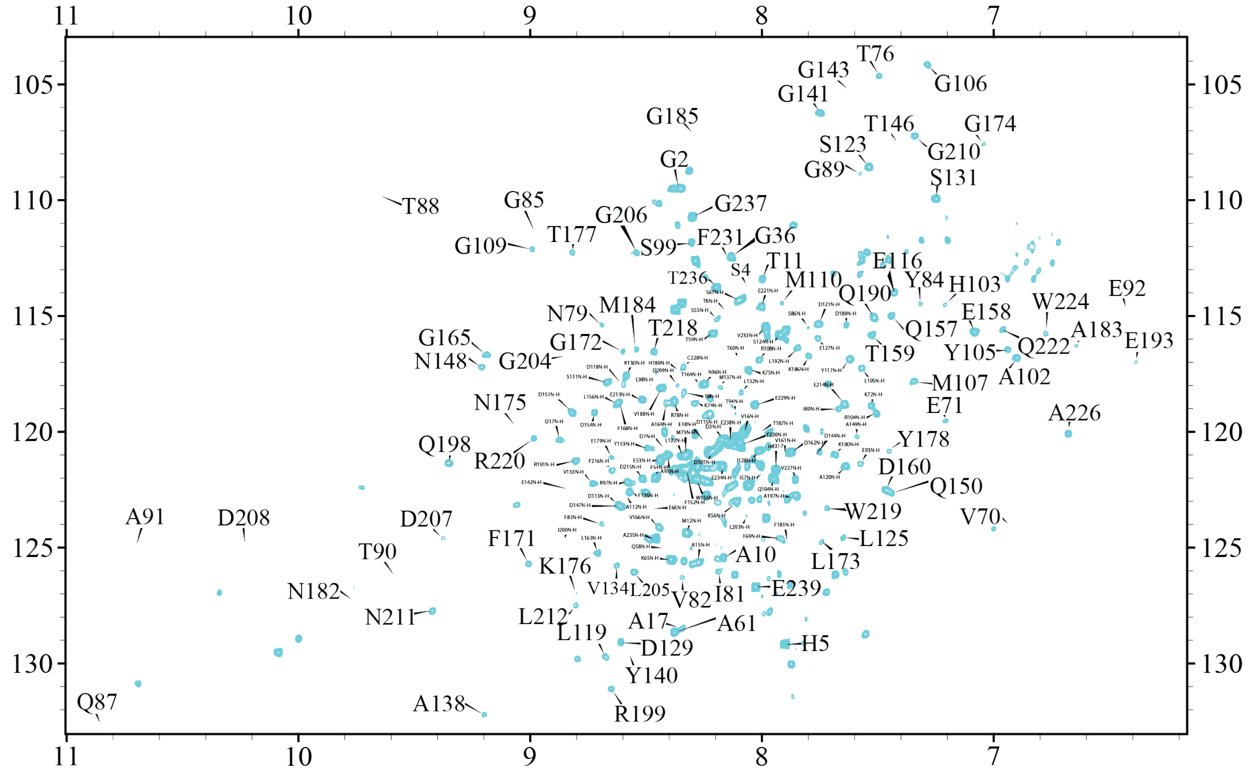


**Figure S18.** A 2D $^{1}H/^{15}N$ TROSY HSQC of $^{15}N$-flFBD in 4F-DMPC ND with 1 molar equivalents of cytP450 and 1-CPI.

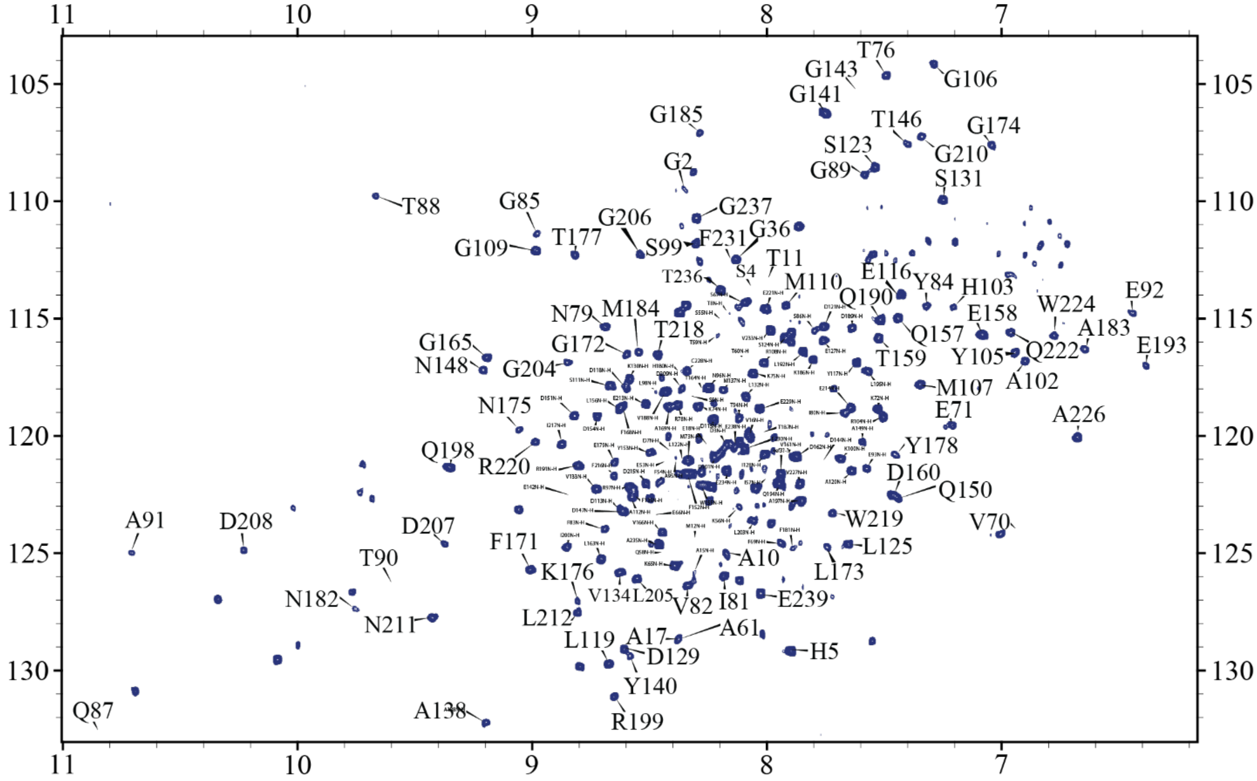


**Figure S19.** A 2D $^{1}H/^{15}N$ TROSY HSQC of $^{15}N$-flFBD in 4F-DMPC ND with 1 molar equivalents of cytP450, 1-CPI, and cytb5.

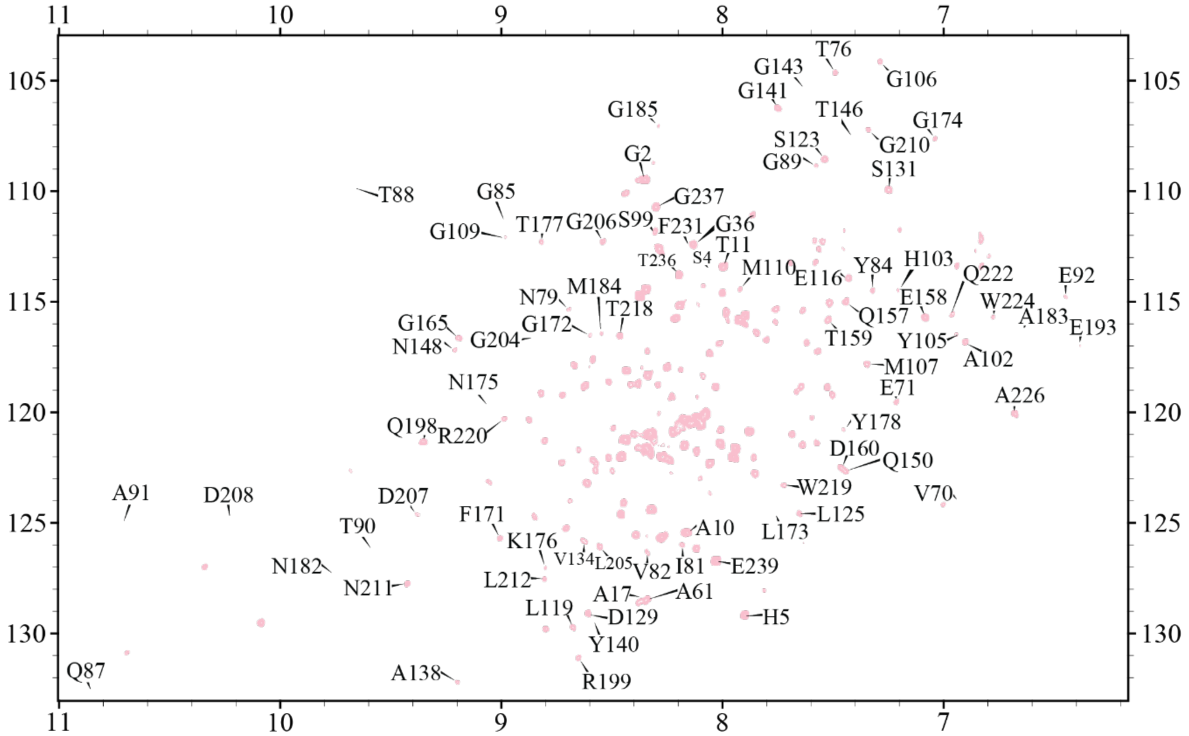


**Figure S20.** A 2D $^{1}H/^{15}N$ TROSY HSQC of $^{15}N$-flFBD in 4F-DMPC ND with 1 molar equivalents of cytP450 and 4-CPI. For clarity, we did not include all of the assignments in this figure, but all were capable of being assigned as in Figure S16.

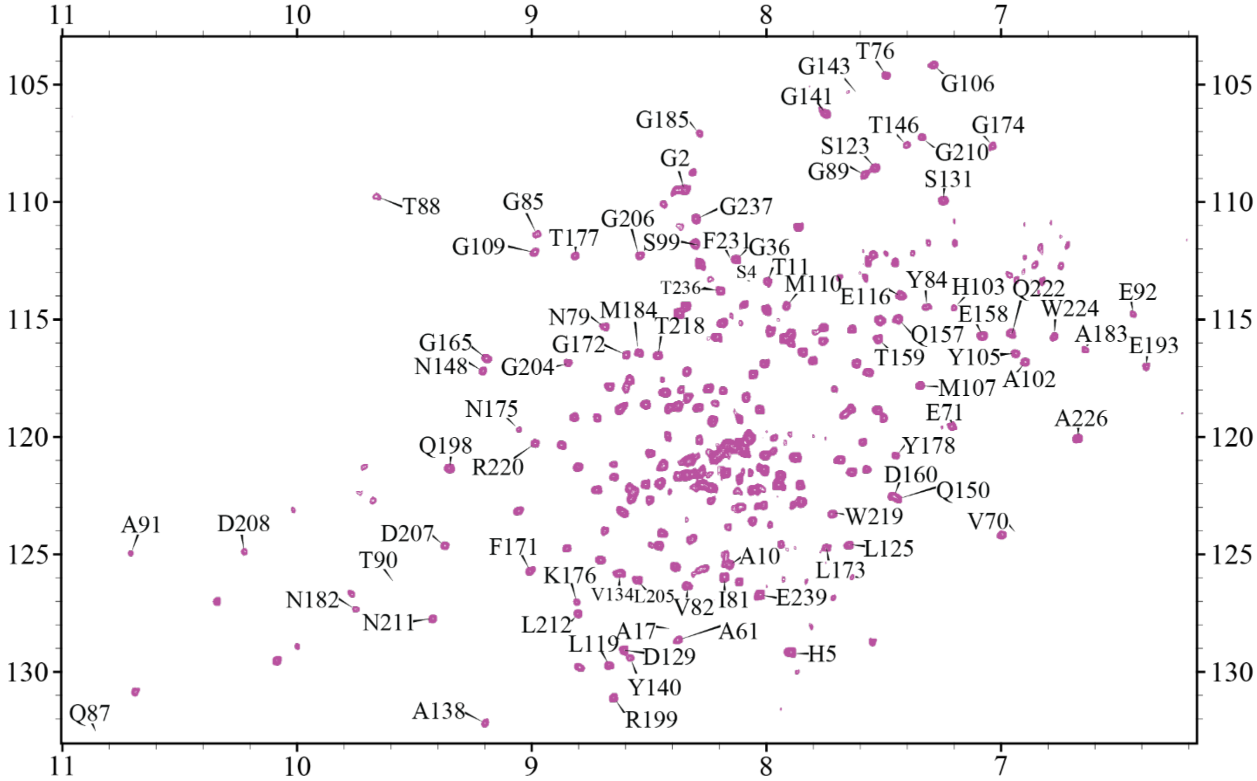


**Figure S21.** A 2D $^{1}H/^{15}N$ TROSY HSQC of $^{15}N$-flFBD in 4F-DMPC ND with 1 molar equivalents of cytP450, 4-CPI, and cytb5. For clarity, we did not include all of the assignments in this figure, but all were capable of being assigned as in Figure S16.

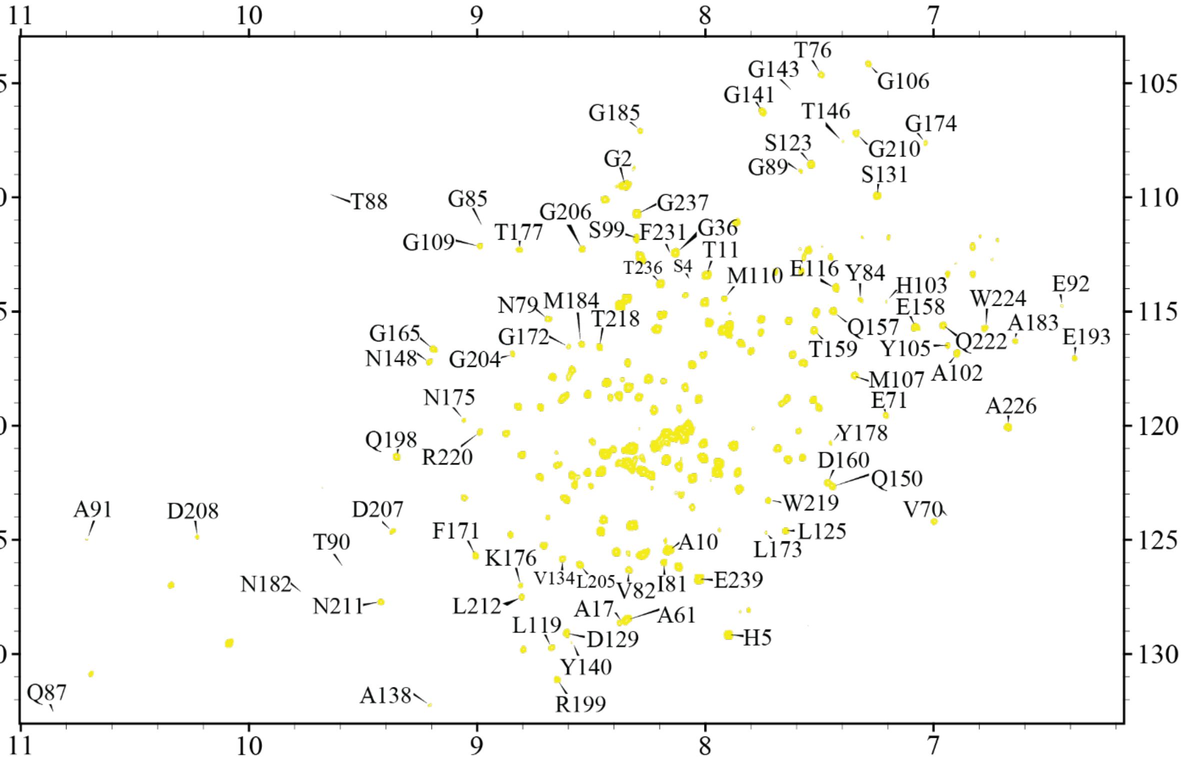


**Figure S22.** A 2D $^{1}H/^{15}N$ TROSY HSQC of $^{15}N$-flFBD in 4F-DMPC ND with 1 molar equivalents of cytP450 and BFZ. For clarity, we did not include all of the assignments in this figure, but all were capable of being assigned as in Figure S16.

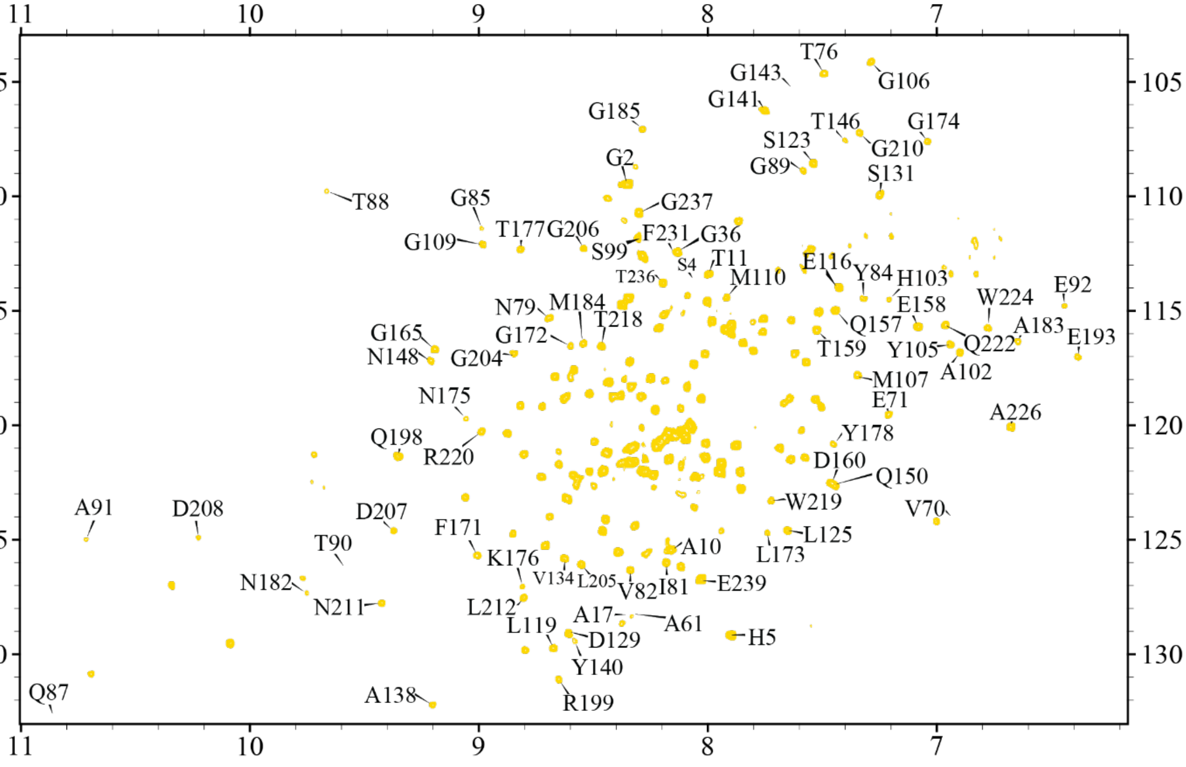


**Figure S23.** A 2D $^{1}H/^{15}N$ TROSY HSQC of $^{15}N$-flFBD in 4F-DMPC ND with 1 molar equivalents of cytP450, BFZ, and cytb5. For clarity, we did not include all of the assignments in this figure, but all were capable of being assigned as in Figure S16.

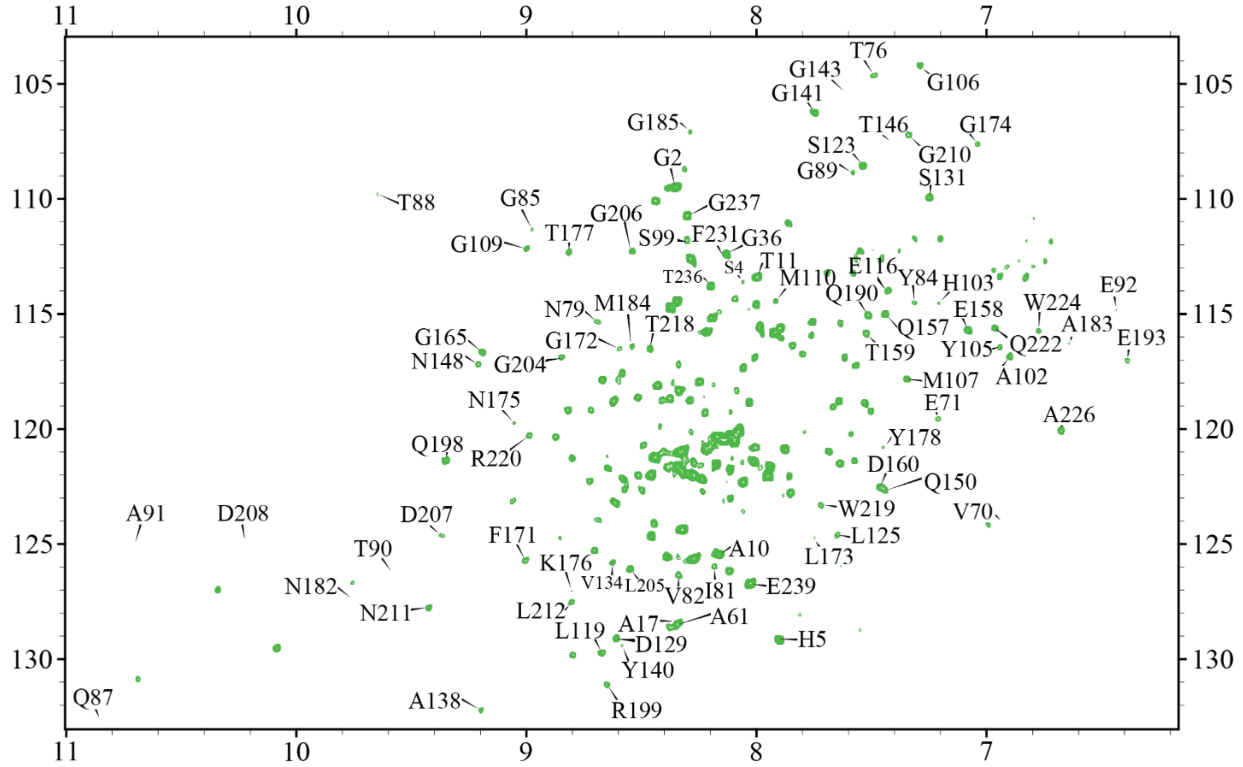


**Figure S24.** A 2D $^{1}H/^{15}N$ TROSY HSQC of $^{15}N$-flFBD in 4F-DMPC ND with 1 molar equivalents of cytP450 and BHT. For clarity, we did not include all of the assignments in this figure, but all were capable of being assigned as in Figure S16.

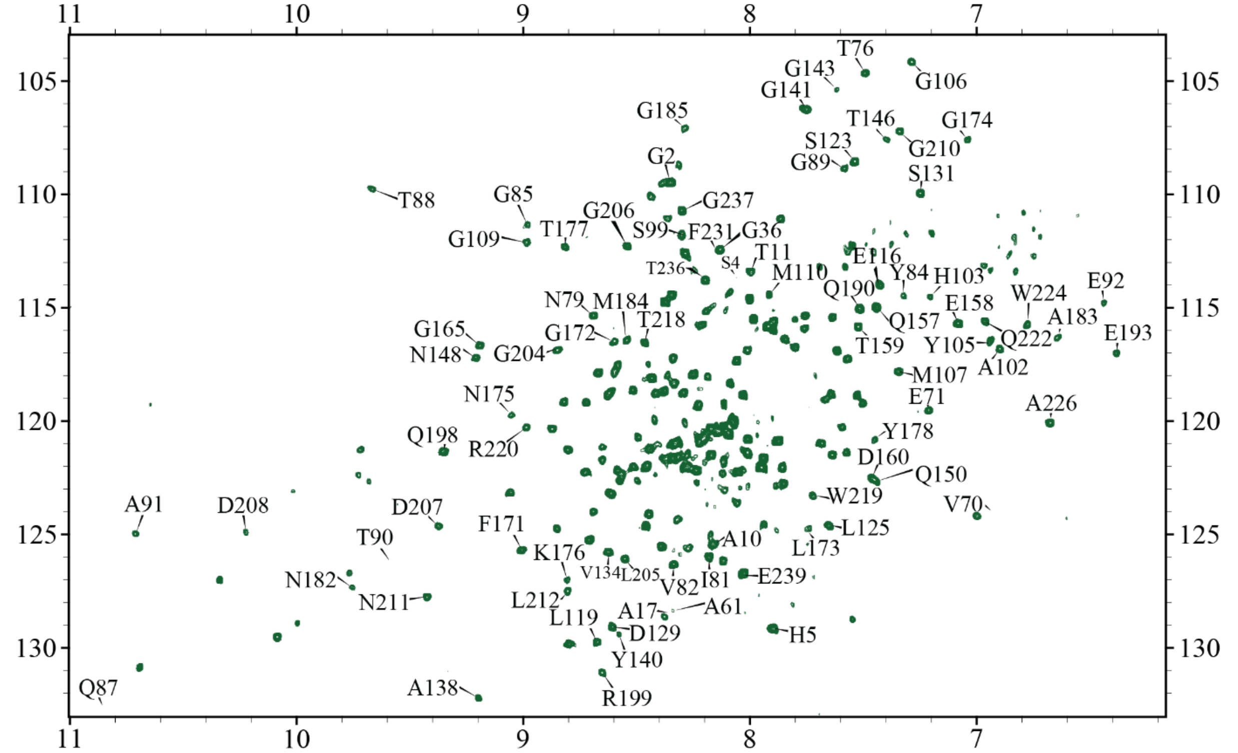


**Figure S25.** A 2D $^{1}H/^{15}N$ TROSY HSQC of $^{15}N$-flFBD in 4F-DMPC ND with 1 molar equivalents of cytP450, BHT, and cytb5. For clarity, we did not include all of the assignments in this figure, but all were capable of being assigned as in Figure S16.

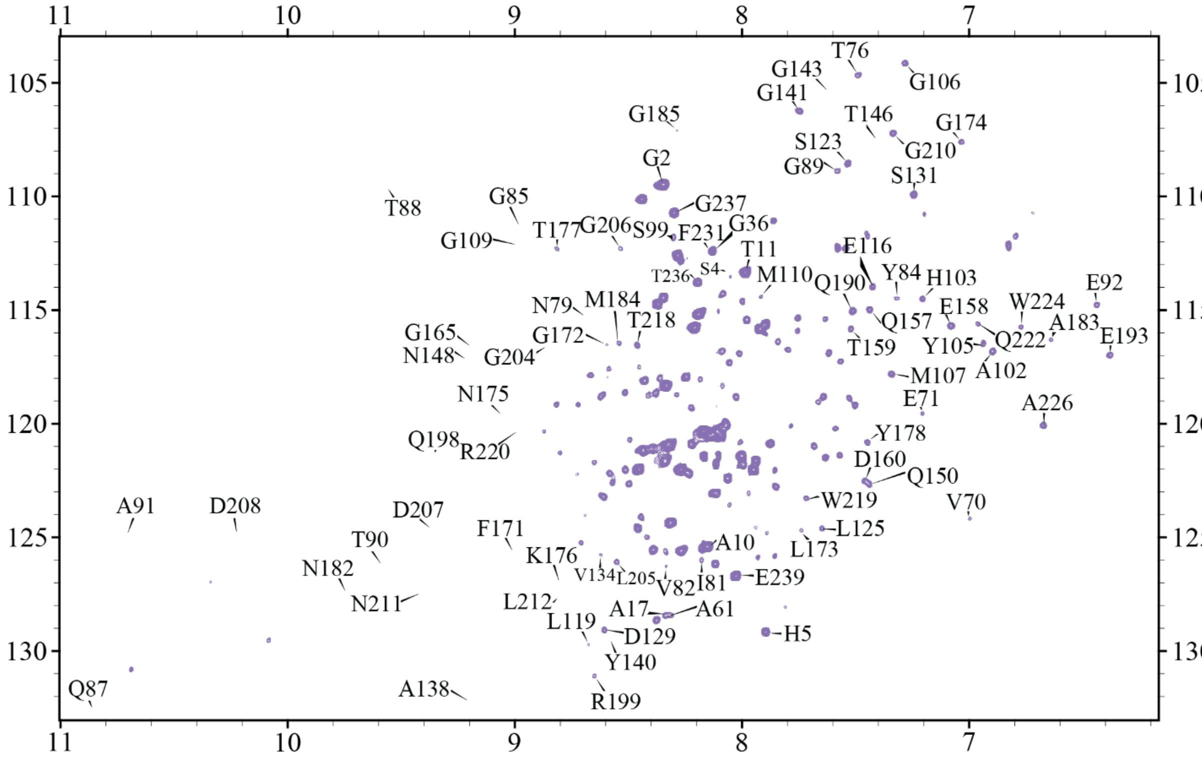


**Figure S26.** A 2D $^{1}H/^{15}N$ TROSY HSQC of $^{15}N$-flFBD in 4F-DMPC ND with 1 molar equivalents of cytP450 and BZ. For clarity, we did not include all of the assignments in this figure, but all were capable of being assigned as in Figure S16.

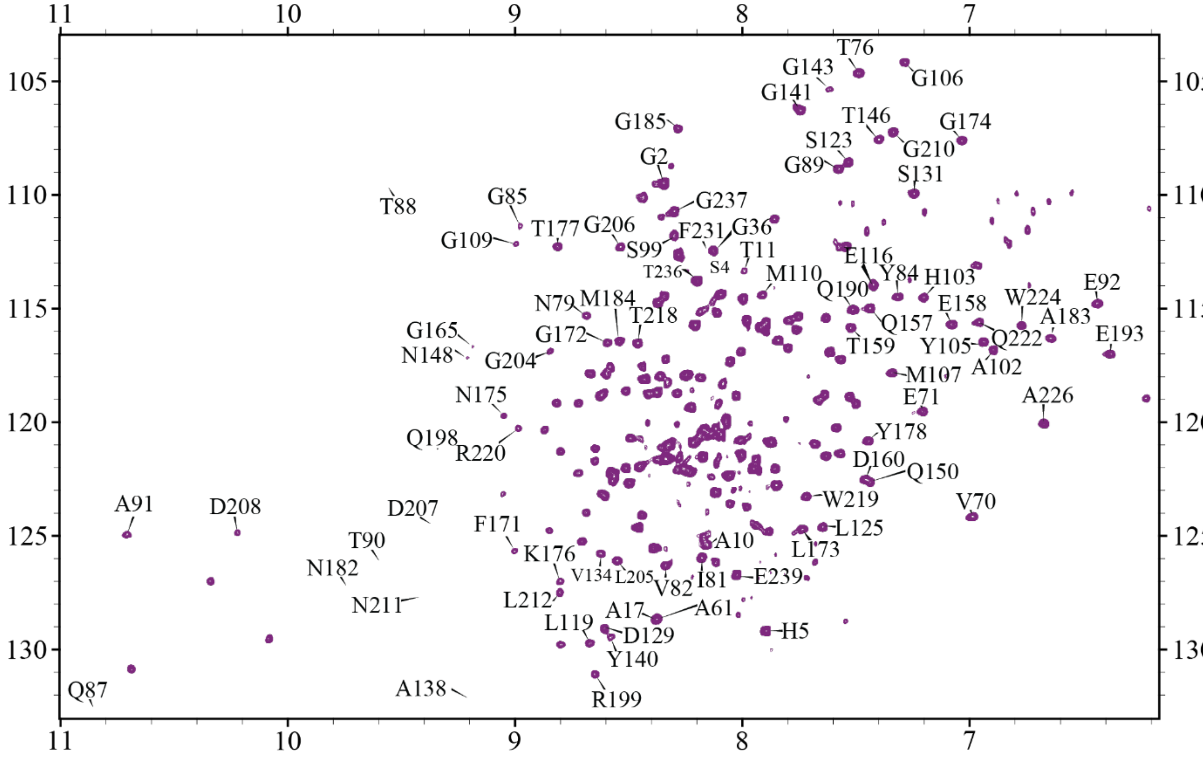


**Figure S27.** A 2D $^{1}H/^{15}N$ TROSY HSQC of $^{15}N$-flFBD in 4F-DMPC ND with 1 molar equivalents of cytP450, BZ, and cytb5. For clarity, we did not include all of the assignments in this figure, but all were capable of being assigned as in Figure S16.

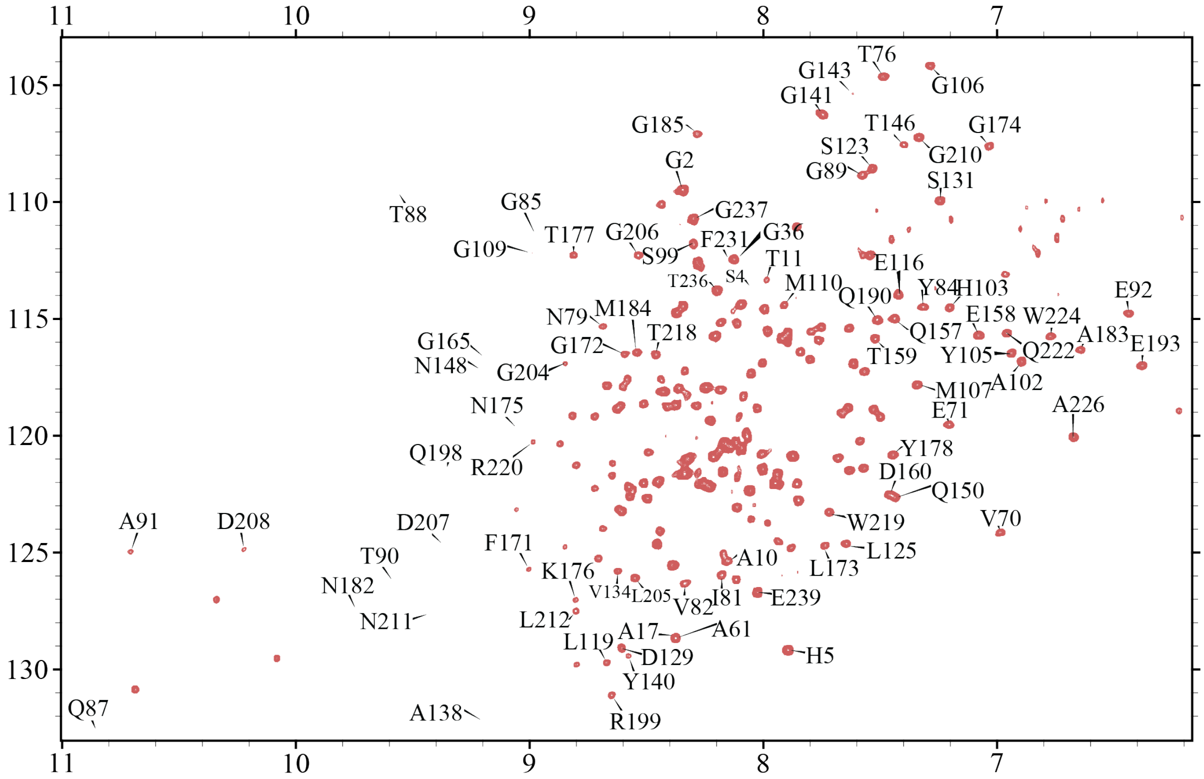


**Figure S28.** A 2D $^{1}H/^{15}N$ TROSY HSQC of $^{15}N$-flFBD in 4F-DMPC ND with 1 molar equivalents of cytP450 and cytb5. For clarity, we did not include all of the assignments in this figure, but all were capable of being assigned as in Figure S16.

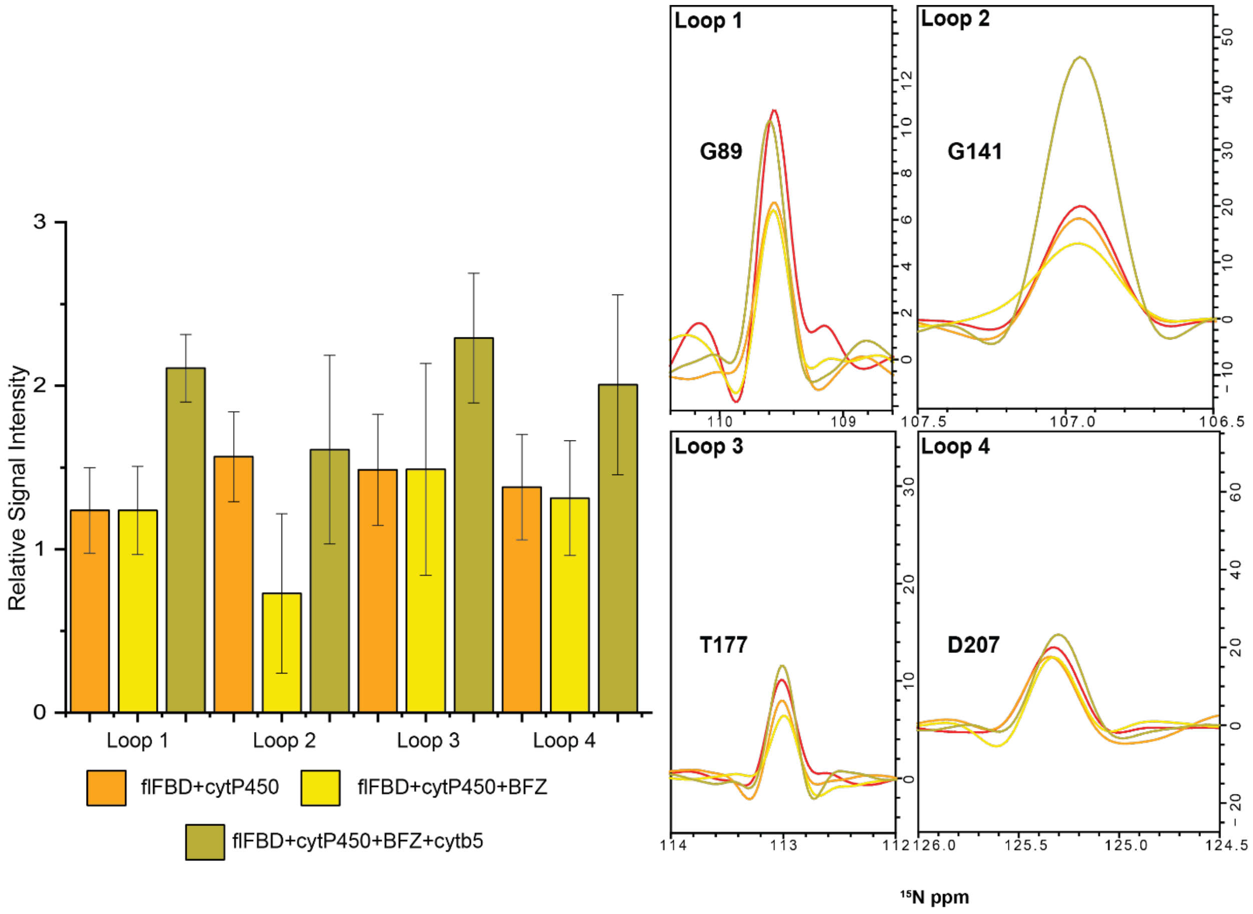


**Figure S29. Loops of flFBD reveal changes in its complexation state.** A. Changes in signal intensities measured from TROSY-HSQC spectra of flFBD for its four loop regions which coordinate its FMN cofactor. Each color represents the different states of flFBD as indicated. B-E. $^{15}$N spectral slices extracted from 2D TROSY-HSQC spectra reveal broadening and restoration of signals for representative peaks of each loop; red trace is for flFBD alone in the ND. The error bars were determined from the standard deviation of the average from the selected residues of each loop (Table S1).

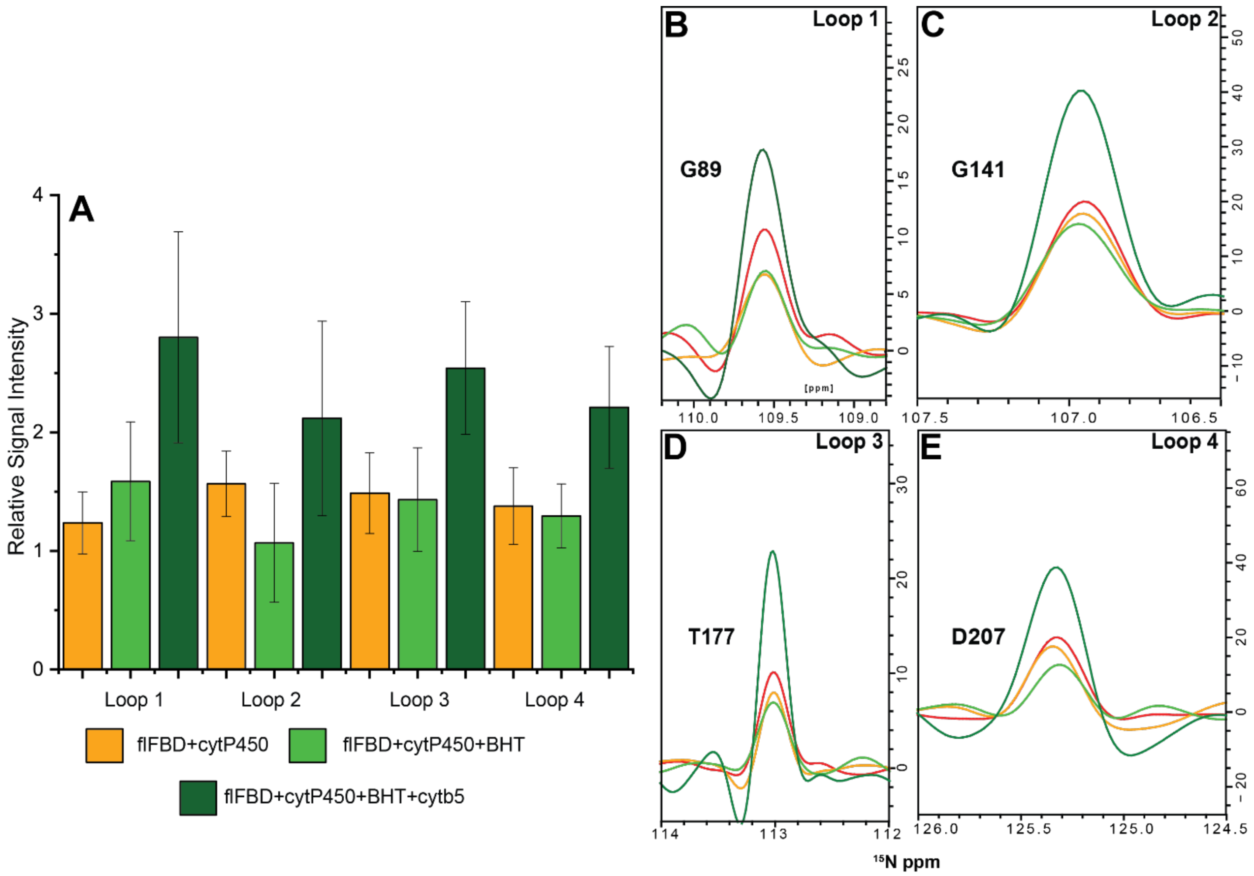


**Figure S30. Loops of flFBD reveal changes in its complexation state.** A. Changes in signal intensities measured from TROSY-HSQC spectra of flFBD for its four loop regions which coordinate its FMN cofactor. Each color represents the different states of flFBD as indicated. B-E. $^{15}N$ spectral slices extracted from 2D TROSY-HSQC spectra reveal broadening and restoration of signals for representative peaks of each loop; red trace is for flFBD alone in the ND. The error bars were determined from the standard deviation of the average from the selected residues of each loop (Table S1).

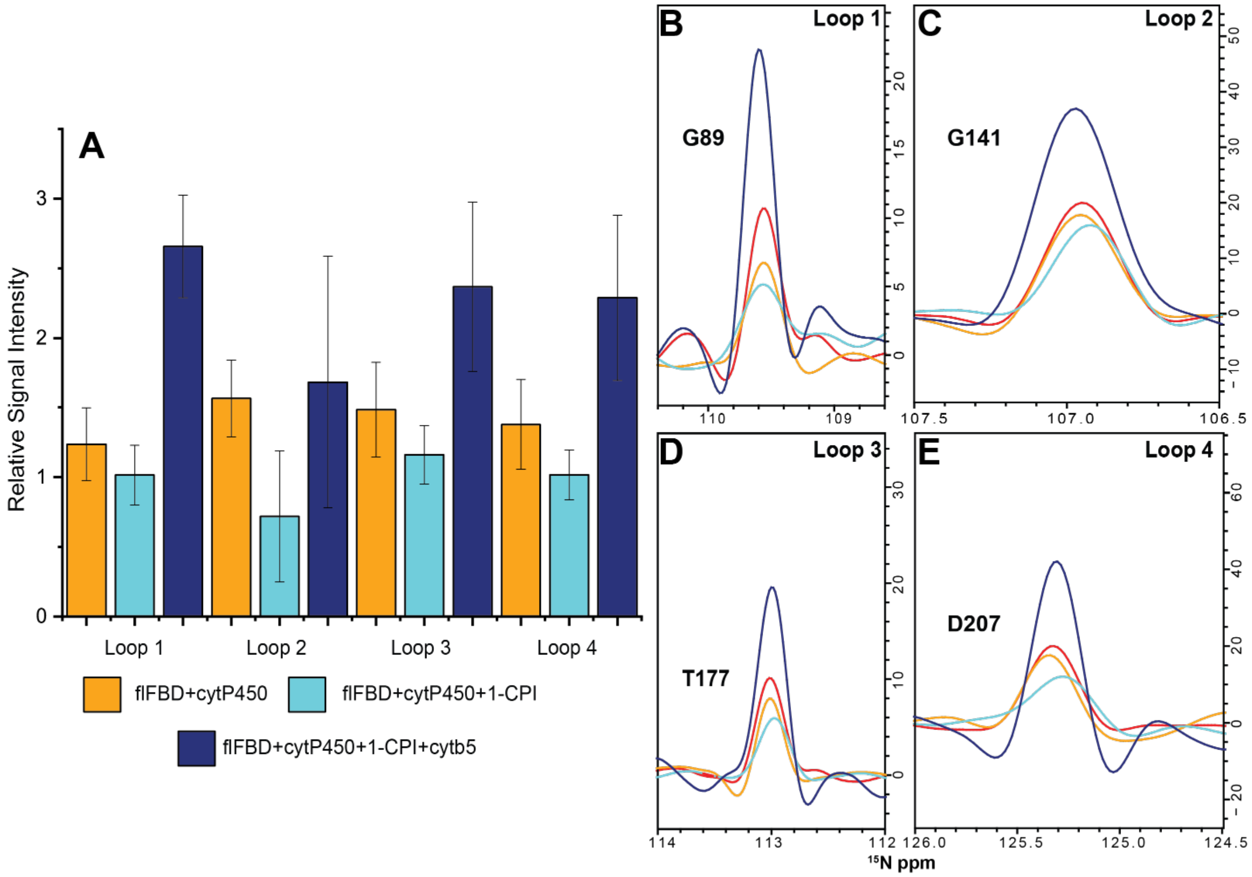


**Figure S31. Loops of flFBD reveal changes in its complexation state.** A. Changes in signal intensities measured from TROSY-HSQC spectra of flFBD for its four loop regions which coordinate its FMN cofactor. Each color represents the different states of flFBD as indicated. B-E. $^{15}$N spectral slices extracted from 2D TROSY-HSQC spectra reveal broadening and restoration of signals for representative peaks of each loop; red trace is for flFBD alone in the ND. The error bars were determined from the standard deviation of the average from the selected residues of each loop (Table S1).

| | Loop Residues |
|---|---|
| Loop 1 | G85, S86, Q87, T88, G89, T90 |
| Loop 2 | Y140, G141, E142, G143, D144 |
| Loop 3 | G174, N175, K176, T177, Y178, E179 |
| Loop 4 | D207, D208, D209, G210, N211, L212 |

**Supplemental Table 1.** A list of the different residues in each of the four loops that coordinate the FMN cofactor in the flFBD.
